# DNA methylation is required to maintain DNA replication timing precision and 3D genome integrity

**DOI:** 10.1101/2020.10.15.338855

**Authors:** Qian Du, Grady C. Smith, Phuc Loi Luu, James M. Ferguson, Nicola J. Armstrong, C. Elizabeth Caldon, Elyssa Campbell, Shalima S. Nair, Elena Zotenko, Cathryn M. Gould, Michael Buckley, Dominik Kaczorowski, Kirston Barton, Ira W. Deveson, Martin A. Smith, Joseph E. Powell, Ksenia Skvortsova, Clare Stirzaker, Joanna Achinger-Kawecka, Susan J. Clark

## Abstract

DNA replication timing and three-dimensional (3D) genome organisation occur across large domains associated with distinct epigenome patterns to functionally compartmentalise genome regulation. However, it is still unclear if alternations in the epigenome, in particular cancer-related DNA hypomethylation, can directly result in alterations to cancer higher order genome architecture. Here, we use Hi-C and single cell Repli-Seq, in the colorectal cancer *DNMT1* and *DNMT3B* DNA methyltransferases double knockout model, to determine the impact of DNA hypomethylation on replication timing and 3D genome organisation. First, we find that the hypomethylated cells show a striking loss of replication timing precision with gain of cell-to-cell replication timing heterogeneity and loss of 3D genome compartmentalisation. Second, hypomethylated regions that undergo a large change in replication timing also show loss of allelic replication timing, including at cancer-related genes. Finally, we observe the formation of broad ectopic H3K4me3-H3K9me3 domains across hypomethylated regions where late replication is maintained, that potentially prevent aberrant transcription and loss of genome organisation after DNA demethylation. Together, our results highlight a previously underappreciated role for DNA methylation in maintenance of 3D genome architecture.

## Introduction

The genome is organised into many higher order architectural layers, including DNA replication timing and 3D chromatin conformation, that serve to functionally compartmentalise genomic regulation. DNA replication follows a highly organised ‘replication timing’ program whereby genomic domains are replicated in a specific temporal order during S-phase, from early to late. The genome is also organised in nuclear space into 3D clusters formed by *cis*-chromatin interactions (Dixon et al., 2012; Lieberman-Aiden et al., 2009; Nora et al., 2012; Sexton et al., 2012). These clusters partition the genome into two large-scale compartments: transcriptionally active, open A-compartments and silenced, mostly closed B-compartments that correspond to the globular architecture of chromatin organisation in interphase nuclei (Lieberman-Aiden et al., 2009). Integration of replication timing (Repli-Seq) and 3D chromatin conformation (Hi-C) sequencing data has revealed that early replication timing domains correspond to the active A-compartments and late replication timing to the repressive B-compartments (Dixon et al., 2012; Ryba et al., 2010). Both DNA replication timing and A/B-compartment structure have been shown to stratify many features of the genome and epigenome. Alterations in DNA replication timing and 3D chromatin organisation correspond to transcriptional and epigenomic changes during differentiation (Miura et al., 2019; Rivera-Mulia et al., 2015) and carcinogenesis (Achinger-Kawecka et al., 2020; Du et al., 2019; Taberlay et al., 2016). These results highlight that DNA replication timing and 3D genome organisation together play a coordinated role in the higher-order regulation of the genome.

One of the major epigenomic hallmarks of cancer is genome-wide loss of DNA methylation (Shen and Laird, 2013), however, it is unknown if hypomethylation directly leads to a deregulation of higher order genome architecture. The DNA methylation landscape of cancer cells has previously been shown to correlate with the large-scale nuclear architecture of the genome, including DNA replication timing landscape and A/B-compartment organisation (Berman et al., 2012; Du et al., 2019; Nothjunge et al., 2017). In particular, long range domains of low DNA methylation called partially methylated domains (PMDs) are associated with late-replicating domains (Berman et al., 2012; Du et al., 2019) and B-compartments (Nothjunge et al., 2017; Xie et al., 2017). These associations raise the question of whether global loss in DNA methylation can directly influence the DNA replication timing program and 3D genome organisation.

Previous loci-specific studies reported that DNA methylation loss was related to changes in replication timing at candidate regions. For example, a shift in replication timing from late to early timing of the inactive chromosome X was associated with DNA demethylation in patients with immunodeficiency disease (ICF) (Hansen et al., 2000) and DNMT1 knockout mouse embryonic stem cells displayed *earlier* replication timing of pericentromeric major satellite repeat elements (Jorgensen et al., 2007). These studies suggest that DNA demethylation can promote alterations in DNA replication timing at a loci-specific level. However, little is also understood about the effect of DNA methylation loss on genome-wide higher order genome organisation.

To investigate the effects of DNA methylation loss on DNA replication timing and 3D genome structure, we used Repli-Seq, single cell Repli-Seq, and Hi-C in a colorectal cancer cell line (HCT116) and its isogenic cell line with double knockout of the maintenance DNA methyltransferase *DNMT1* and *de novo* DNA methyltransferase *DNMT3B* (DKO1) (Rhee et al., 2002). We show that hypomethylation, a common hallmark of cancer, results in loss of replication timing precision and concordant deregulation of 3D genome organisation. Our results highlight a previously underappreciated role for DNA methylation in the direct regulation of the 3D genome.

## Results

### Co-ordinate alterations to DNA replication timing and 3D genome organisation

To directly determine the impact of DNA methylation levels on higher order genome architecture, we investigated changes in both DNA replication timing and 3D chromatin conformation after global DNA hypomethylation. We used a well-described model of DNA methylation loss; HCT116, a colorectal cancer cell line and DKO1, the isogenic cell line with double knockout of the maintenance DNA methyltransferase *DNMT1* and the *de novo* DNA methyltransferase *DNMT3B* (Rhee et al., 2002). We found that the DKO1 cells show genome-wide DNA hypomethylation (~56% methylation loss) compared to HCT116 cells (**Fig. 1a**). To determine if global DNA methylation loss results in changes in DNA replication timing and/or 3D genome structure, we performed Repli-Seq and *in situ* Hi-C in duplicate in HCT116 and DKO1 cells (**See Methods, Supp Fig. 1, Supp Fig. 2a**).

**Figure 1:**
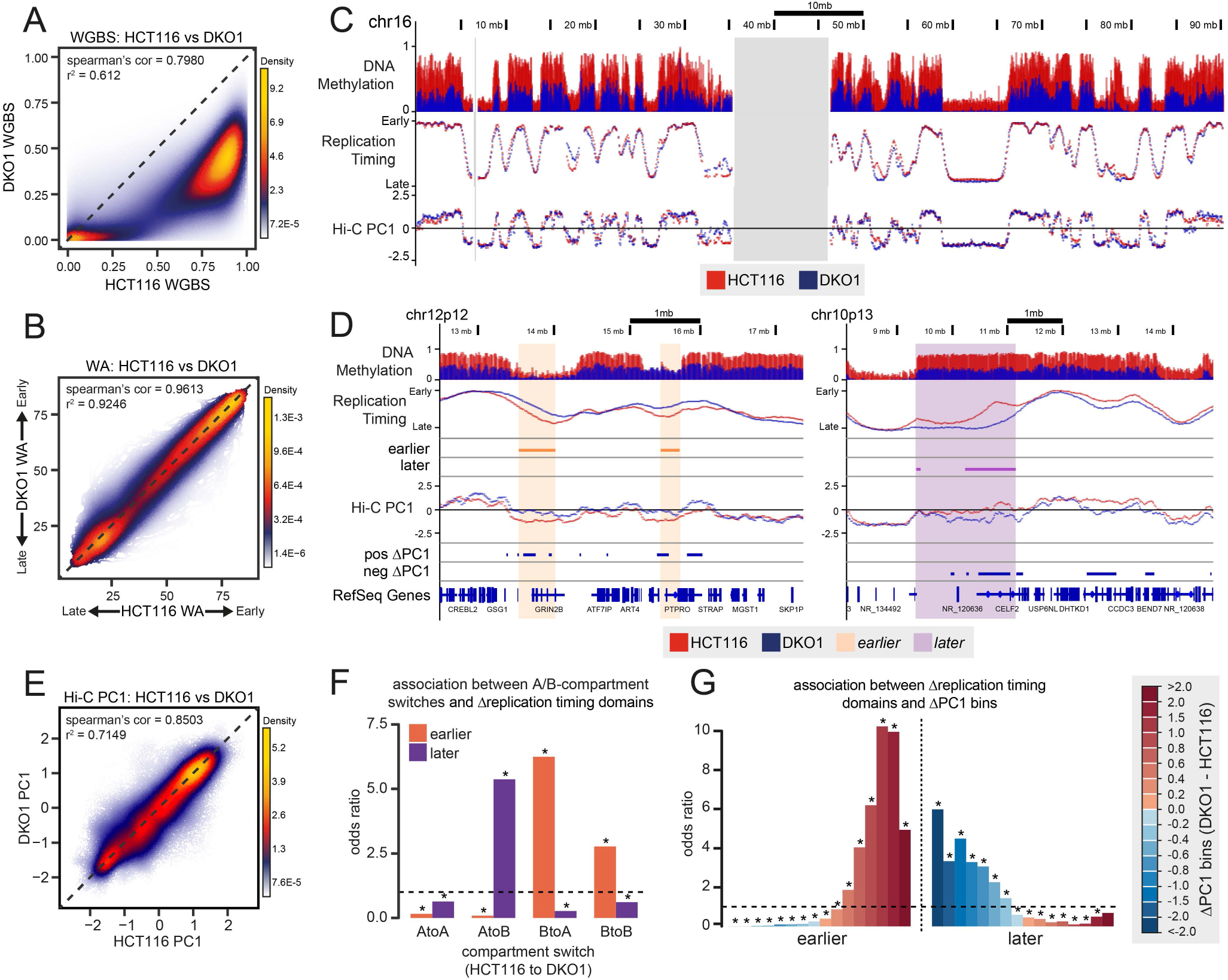
Co-ordinate change in nuclear organisation and DNA replication timing following DNA methylation loss. **A** Heat scatterplot showing replicate averaged methylation levels of HCT116 against averaged methylation levels of DKO1. **B** Heat scatterplot showing replicate averaged replication timing values (WA) of HCT116 against averaged replication timing values of DKO1. **C** Representative examples of replication timing and Hi-C PC1 profiles in HCT116 (red) and DKO1 (blue). The grey bars indicate regions of the genome where there is no data, in this case the centromeric region. **D** Representative examples of regions showing concordant change in replication timing and Hi-C PC1. **E** Heat scatterplot showing replicate averaged Hi-C PC1 values of HCT116 against DKO1. **F** Fisher’s exact test for association between A-/B-compartment switches and *earlier/later* replicating loci. **G** Fisher’s exact test for association between intervals of ΔPC1 to *earlier* and *later* replicating loci. For **F** and **G**, asterisks indicate significant associations (FDR < 0.05) and dotted line indicates an odds ratio of 1.

We first examined genome-wide trends in DNA replication timing and Hi-C data between HCT116 and DKO1. We found that in contrast to global DNA hypomethylation (**Fig. 1a**), replication timing weighted average (WA) values between HCT116 and DKO1 are highly correlated (**Fig. 1b**, Spearman’s = 0.9613). A representative example is shown in **Fig. 1c**. We next used a WA difference of 15 (|ΔWA| >15), as a stringent approach to define regions with large replication timing alterations, and a WA difference of 5 (|ΔWA| < 5) to measure minimal changes in timing (**See Methods**, **Supp Fig. 1e & f**). We found that 66.8% of the genome shows close conservation of replication timing (|ΔWA| < 5) and 29.84% of the genome displays a moderate shift in replication timing (5 < |ΔWA| < 15). Notably, at the most stringent cut off (|ΔWA| >15), we found a distinct fraction of the genome displayed a large shift in replication timing, that is to either replicate *earlier* (1.89%) or replicate *later (*1.47%) in DKO1 compared to HCT116 (**Fig. 1d**).

Next, to investigate if loss of DNA methylation results in alterations to large-scale genome compartmentalisation, we performed compartment analyses to define A/B compartment switching from HCT116 to DKO1. We defined A/B compartment status with the first principal component (PC1) values, which represent euchromatin/heterochromatin neighbourhoods, respectively. Similarly to replication timing, we observe that HCT116 and DKO1 Hi-C data show high correlation between PC1 values (**Fig. 1c,e**, **Supp Fig. 2b**) and comparable proportions of A and B compartments (**Supp Fig. 2c**). We found that 13.58% of the genome had switched A/B compartments comprising switching from A to B (9.10%) and switching B to A (4.48%) (**Supp Fig. 2d**). A large proportion of compartment switches were centred around PC1 values of <0.5 and > PC1 > −0.5 (**Supp Fig. 2e**), indicating that, while compartment switches do cross the midline that defines A vs. B compartments, a large proportion of these switches are at regions with low compartment values (**Supp Fig. 2f**). Therefore, to identify regions which show strong compartment shifts, regardless of whether they cross the midline, we defined switching regions by their ΔPC1 score, using a ΔPC1 cut-off of ≥ |1| (**Fig. 1d**, **Supp Fig. 2g**, **See Methods**). Using these criteria, we found 4.43% of the genome have increased PC1 values in DKO1 and therefore is more A-type than HCT116, and 3.47% of the genome have decreased PC1 values where DKO1 is more B-type than HCT116.

Lastly, we examined whether there is concordance between regions that show a change in genome compartmentalisation and regions that show changes in DNA replication timing, following DNA hypomethylation. We found that domains that replicated *earlier* are enriched for B to A compartment switches and positive ΔPC1 changes, and *later* replicating domains are enriched for A to B compartment switches and negative ΔPC1 changes (**Fig. 1f,g**). As expected, where DNA replication timing has become *earlier* in DKO1 compared to HCT116, the region moves more towards the ‘active’ A compartment (positive PC1 values) and vice versa (**Fig. 1f,g**). Exemplary regions are shown in **Fig. 1d**. In summary, we found that in response to global DNA hypomethylation, ~3-10% of the genome undergo large and co-ordinate changes in higher-order genome architecture.

### Changes in higher order genome architecture occur at partially methylated domain boundaries

We next examined whether the ~3-10% of the genome that show large changes in replication timing and genome organisation are related to the degree of methylation loss. Surprisingly, we found that loci that replicated *earlier* in DKO1 are significantly associated with a gain in methylation (**Fig. 2a**). Similarly, we found that regions with a large positive ΔPC1 difference from HCT116 to DKO1 are also enriched for regions that predominately show a gain in methylation (**Fig. 2b**). Whereas loci with DNA methylation loss are slightly associated with loci that replicated *later* or showed a large negative ΔPC1 difference from HCT116 to DKO1 (**Fig. 2a,b**). This suggests that the distinct higher order genome architectural changes may occur co-ordinately with the degree of DNA methylation change, however, appear to be more related to methylation gain rather than methylation loss. As the DNA methylation landscape in HCT116 cells is uneven across DNA replication timing (**Fig. 1c**), different replication times are subject to different amounts of methylation change. Early replicating loci are highly methylated and are thus able to lose methylation, whilst late replicating loci are lowly methylated and thus more amenable to methylation gain. Loci that show methylation gain are indeed lowly methylated in HCT116 and conversely loci that show methylation loss have high methylation in HCT116 (**Supp Fig. 3a**). In agreement, we found that *earlier* replicating and positive ΔPC1 loci that are associated with methylation gain are lowly methylated and *later* replicating and negative ΔPC1 loci that associate with methylation loss are highly methylated (**Supp Fig. 3b**).

**Figure 2:**
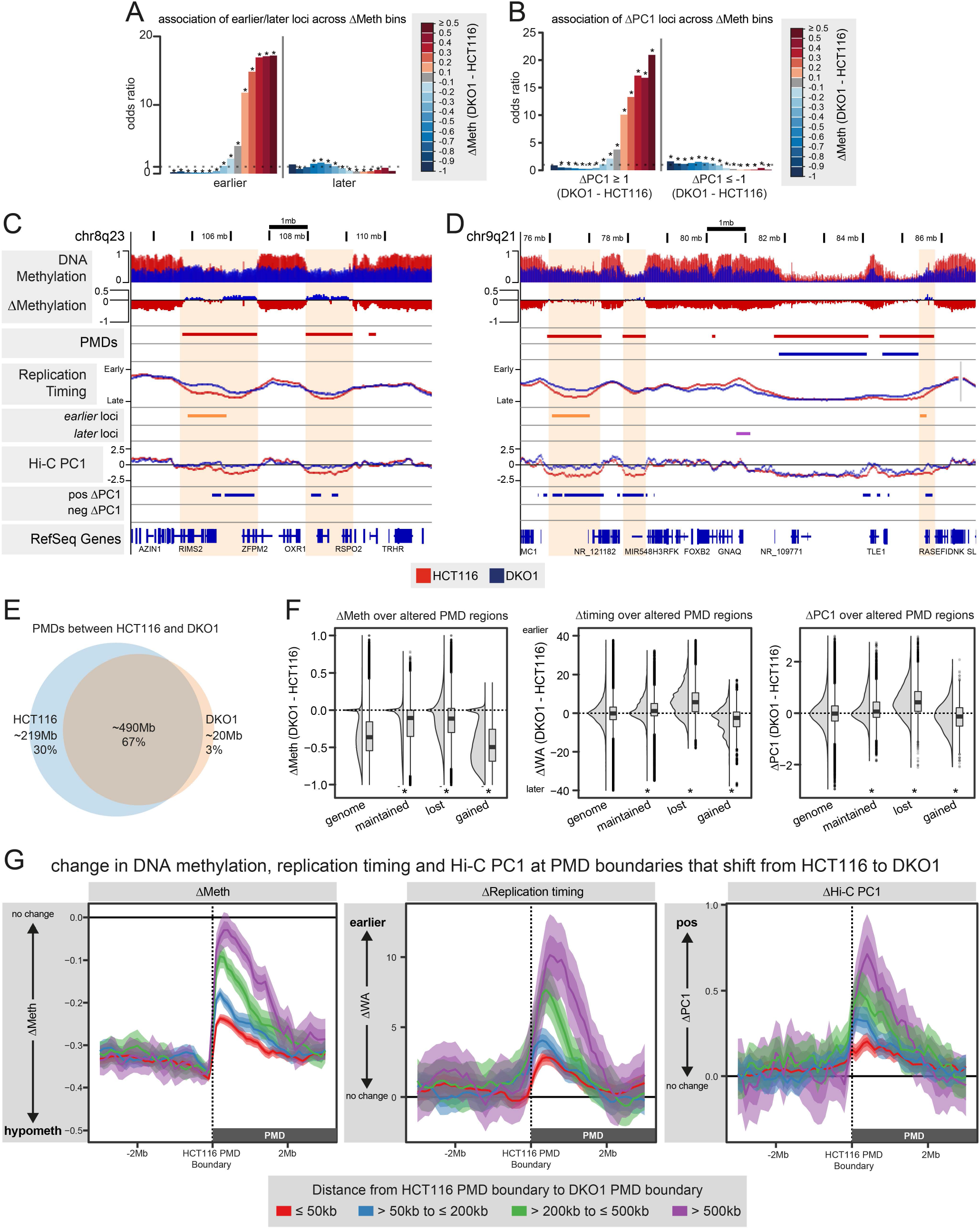
Changes in replication timing and nuclear organisation occur at partially methylated domain boundaries. **A** Fisher’s exact test for association between *earlier* and *later* loci across bins of methylation change. **B** Fisher’s exact test for association of domains of ΔPC1 across bins of methylation change. For **A** and **B**, asterisks indicate significant associations (FDR < 0.05) and dotted line indicates an odds ratio of 1. **C,D** Representative examples of regions containing lost PMDs. **E** Overlap of PMDs between HCT116 and DKO1. **F** The change in methylation, replication timing and Hi-C PC1 at maintained, lost or gained PMDs compared to genome-wide ‘genome’. Asterisks indicate significance (*p* < 0.05) in one-tailed Mann-Whitney-Wilcoxon test against ‘genome’. For maintained and lost, alternative is ‘greater’. For gained, alternative is ‘less’. **G** HCT116 PMDs are grouped into 4 groups based on the distance of shift inwards of the DKO1 PMD boundary from the HCT116 PMD boundary at the 5’ and/or 3’ end. Profile plots of the average change in DNA methylation, change in replication timing and change in Hi-C PC1 of the four groups are shown. Plots show an average line with width of shading indicating confidence intervals.

Visual inspection of DNA methylation, replication timing and compartment structure revealed that changes in replication timing and chromatin conformation frequently co-occur at troughs in the DNA methylation profile, also known as partially methylated domains (PMDs) (**Fig. 2c,d** and **Supp Fig. 3c**). PMDs are a characteristic feature of the cancer DNA methylation landscape (Berman et al., 2012; Hon et al., 2012; Hovestadt et al., 2014) and correspond to late replication domains (Berman et al., 2012) as well as B compartments (Nothjunge et al., 2017; Xie et al., 2017).

Interestingly, we found that despite widespread hypomethylation in DKO1 cells, the majority of HCT116 PMD regions persist in DKO1 (**Fig. 2e**). Both HCT116 and DKO1 PMDs are late-replicating and form ‘troughs’ in the DNA methylation profile, replication timing profile and PC1 compartment values (**Supp Fig. 3d**). However, DKO1 PMDs have less well-defined DNA methylation boundaries due to the global hypomethylation (**Supp Fig. 3d**) and also show shallower ‘troughs’ in both the replication timing and Hi-C PC1 profiles.

To explore the alterations in PMD structure, we divided PMDs into regions that are maintained, lost or gained from HCT116 to DKO1 (**See Methods**). Interestingly, regions that lose PMD definition associate with the same bins of methylation change as *earlier* replicating and positive ΔPC1 compartment regions (**Supp Fig. 3e**). Furthermore, lost PMD regions show a clear shift towards early replication timing and positive PC1 values in DKO1 compared to HCT116 (**Fig. 2f**), contributing to ~50% of all *earlier* loci and ~34% of all positive ΔPC1 regions (**Supp Fig. 3f**). We further observed that PMD loss appears to occur specifically at the boundaries of the HCT116 PMD and shifts inwards from HCT116 to DKO1 (**Fig. 2d**, **Supp Fig. 3c**). We found that larger shifts in PMD boundaries show larger changes towards *earlier* replication timing, larger increases in Hi-C PC1 values (i.e. B to A shift) and less methylation loss between HCT116 and DKO1 (**Fig. 2g**). Therefore, the shift in PMD boundaries, due to the loss of the stratification of the DNA methylation landscape, is associated with a corresponding shift in higher-order genome architecture at PMD boundaries.

### DNA hypomethylation reduces precision of DNA replication timing

Interestingly, even though replication timing values between HCT116 and DKO1 are highly correlated (**Fig. 1b**, Spearman’s = 0.9613), we observed that the range of pre-normalised replication timing (WA) values in DKO1 cells is smaller compared to HCT116 (**Supplementary Fig. 1d**). In the Repli-Seq method, we calculate WA values from 6-fraction values across the S-phase called percentage-normalised density values (PNDVs) (**See Methods**). To investigate the difference in spread of WA values between HCT116 and DKO1, we first determined the variance of the 6-fraction signal. A higher score indicates that the majority of the signal is coming from a small number of fractions (precise timing), whereas a lower score indicates the signal is more evenly distributed between the 6-fractions (varied timing) (**Supp Fig. 4a**). We found that DKO1 showed lower variance than HCT116 (**Fig. 3a**), suggesting that replication in DKO1 is more spread over the 6-fractions and therefore less precise. Representative examples of regions with decreased precision in DKO1 compared to HCT116 are shown in **Supplementary Fig. 4b**.

**Figure 3:**
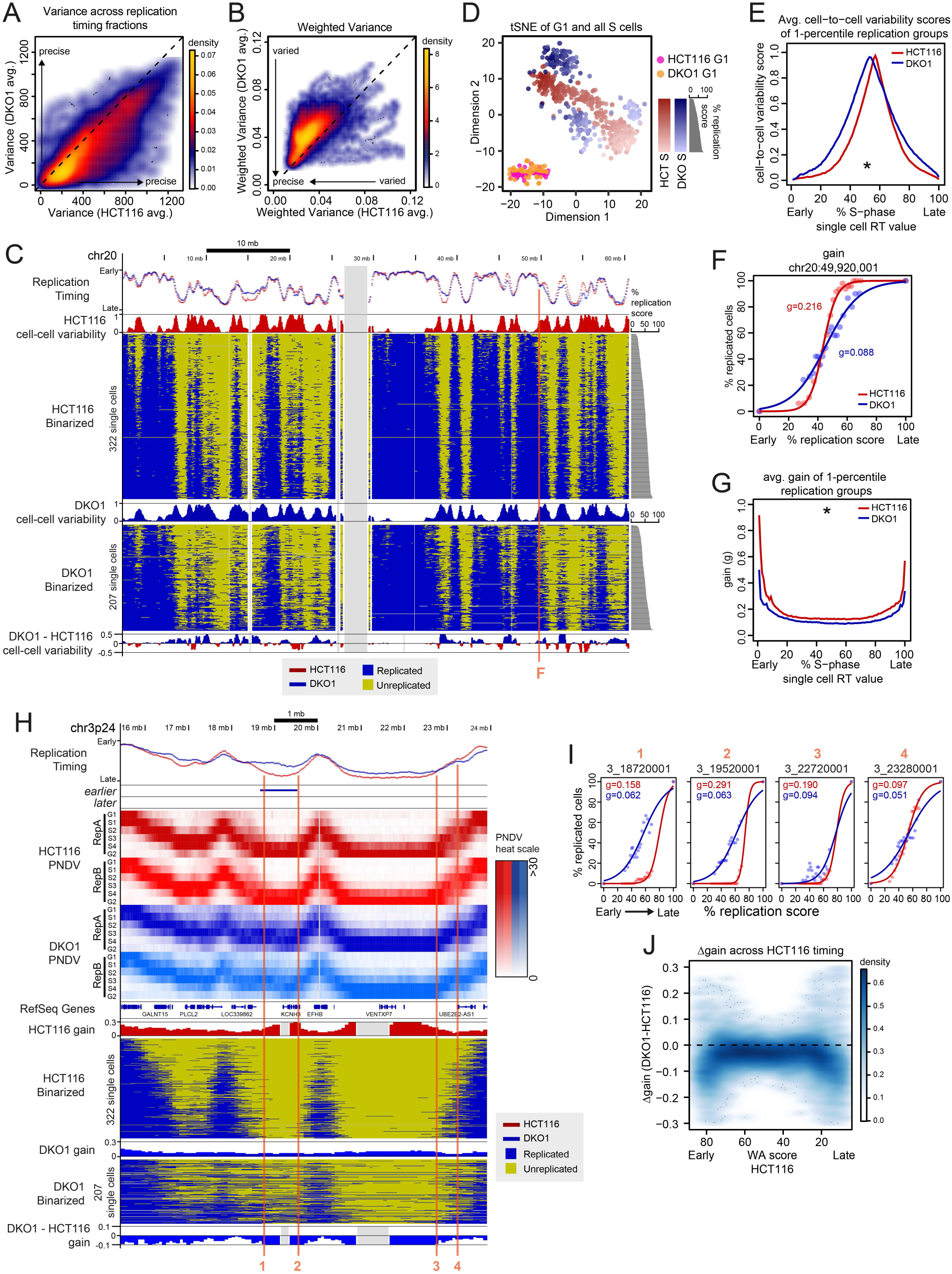
Loss of global methylation increases cell-to-cell heterogeneity of DNA replication timing. **A** Heat scatterplot of average HCT116 variance scores against average DKO1 variance scores. As indicated, the higher the variance value, the more precise the replication timing score. **B** Heat scatterplot of average HCT116 weighted variance scores against average DKO1 weighted variance scores. As indicated, the lower the weighted variance value, the more precise is the replication timing score. **C** Representative example of whole population Repli-Seq with single cell Repli-Seq (scRepli-Seq). Binarised scRepli-Seq data is ordered from earliest (lowest % replication score) to latest (highest % replication score) cells. Also shown are cell-to-cell variability scores for each cell line and the difference. Red vertical line refers to the 80kb bin shown in **F**. The grey bars indicate regions of the genome where there is no data, in this case the centromeric region. **D** tSNE of G1 and S-phase cells from HCT116 and DKO1. Early to late gradient of S-phase cells is denoted by transition from dark to light shading. **E** Average cell-to-cell variability score per 1-percentile of bins across single cell RT values. Asterisk indicates significant difference (*p* < 0.05) between HCT116 and DKO1, calculated using a permutation test. **F** Sigmoid model curves and gain values of example locus from **C**. Dots are real data points and lines are the fitted curves. **G** Average gain scores per 1-percentile of bins across single cell RT values. Asterisk indicates significant difference (*p* < 0.05) between HCT116 and DKO1, calculated using a permutation test. **H** Representative region with increased heterogeneity (loss of gain value) in DKO1 compared to HCT116. All 6 PNDV fractions for each whole population Repli-Seq dataset is shown in the order of G1, S1, S2, S3, S4 and G2 from top to bottom. PNDV value for each fraction is indicated by a heat colour scale. Binarised scRepli-seq data is ordered from earliest (lowest % replication score) to latest (highest % replication score) cells. Gain scores are shown for each cell line and the difference. Red vertical line refers to 80kb bins shown in **I**. **I** Sigmoid model curves and gain values of loci from **H** that showed increased heterogeneity in DKO1 compared to HCT116. Dots are real data points and lines are the fitted curves. **J** Heat scatterplot of gain differences (DKO1 – HCT116) across whole population replication timing values (WA) of HCT116.

A standard variance calculation disregards the order of the 6 fractions, whereas the order of the replication timing fractions across S-phase is biologically important. Therefore, we re-examined the variation using a weighted variance calculation (**See Methods**). Here, a lower value indicates more precision in timing (**Supp Fig. 4e**). Using the weighted variance score, we again showed that the majority of loci show more variance and reduced precision of DNA replication timing in DKO1 compared to HCT116 (**Fig. 3b**). This data suggests that there is a potential decrease in the synchronisation of DNA replication timing, resulting in a decrease in replication timing precision that occurs as a result of DNA hypomethylation.

### DNA hypomethylation increases cell-to-cell heterogeneity of DNA replication timing

To determine if the reduction in the precision of population-level Repli-Seq data is due to an increase in cell-to-cell variability or heterogeneity within the cell population, we performed single cell replication timing sequencing (scRepli-Seq) (Takahashi et al., 2019). Single cell libraries were generated on the Single Cell CNV Solution platform from 10x Genomics (**See Methods**, **Fig. 3c,d**). Single G1 and S-phase cells cluster within their respective cell cycle states between HCT116 and DKO1 (**Fig. 3d**, **Supp. Fig 4f**). Early S-phase cells are closer to the G1 population and late S-phase cells are further from the G1 population along dimension 1 (**Fig. 3d**). In line with lack of replication timing precision, DKO1 cells are more disparately distributed within the S-phase cell cluster relative to HCT116 cells. We next calculated cell-to-cell variability using mid-S-phase cells (40-70% replication) for 80kb bins across the genome. Cell-to-cell variability scores are highest at mid-replication timing (**Supp Fig. 4g**), similar to Takahashi *et al.* (2019). We found that average cell-to-cell variability scores of loci in one-percentile groups across replication timing were statistically higher in DKO1 compared to HCT116 (**Fig. 3e**), confirming increased cell-to-cell variability in the hypomethylated DKO1 cells.

Next, to address if the degree of replication heterogeneity throughout S-phase is also higher in DKO1 cells compared to HCT116 cells, we performed sigmoidal curve modelling of the single cell data to obtain replication kinetics for 80kb bins across the genome (**See Methods**). An exemplary region from **Fig. 3c** with sigmoidal curve modelling is shown in **Fig. 3f**. The gain (or slope) of each sigmoid curve indicates how heterogeneously a locus is replicated between cells. A steep curve (large gain value in HCT116, e.g. g=0.216) indicates synchronous replication of the loci amongst cells, and a flatter curve (small gain value in DKO, e.g. g=0.088) indicates heterogeneous replication of the loci amongst cells. Similar to Takahashi et al. (2019), gain value is highest towards the earliest and latest extremes of replication timing, indicating start and end of replication has the least cell-to-cell heterogeneity (**Supp Fig. 4h**). Again, DKO1 cells showed overall lower gain values genome-wide than HCT116 across replication timing, indicating more heterogeneous replication within the DKO1 cell population (**Fig. 3g**). Representative examples of the consistent lower gain values in DKO cells can be seen in **Fig. 3h** (lower panel DKO1-HCT116) and **Fig. 3i**. This trend in increased heterogeneity occurs genome-wide, including regions with (**Fig. 3i loci 1,2**) and without a change in replication timing (**Fig. 3i loci 3,4, Supp. Fig. 4i**), and particularly at very early and very late regions (**Fig. 3j**). Increase in heterogeneity at the beginning and end of S-phase agrees with the reduced range of population-level Repli-Seq WA values in DKO1 (**Supp. Fig. 1d**) and suggests DNA hypomethylation has caused an ‘erosion’ in the precise regulation of replication timing.

### DNA hypomethylation reduces integrity of 3D genome compartmentalisation

We next examined the effects of reduced replication timing precision on 3D genome organisation. As Hi-C is a cell population-level assay, we hypothesise that similar to loss of the precision of replication timing, we may observe a loss of strength of organisation. In corroboration with replication timing cell-to-cell heterogeneity, we found that DKO1 cells have reduced compartmentalisation strength of both A and B compartments, indicated by the reduction of intra-compartment contact frequencies (A-A, B-B) and an increase of inter-compartmental (A-B) contact frequency (**Fig. 4a,b**). However, DKO1 have become less B-B interaction dominant overall (**Fig. 4c**), suggesting that B compartments are more affected by global DNA hypomethylation. We also observe loci that have maintained B-compartment status are enriched in *earlier* replicating regions (**Fig. 1f**), supporting that ‘inactive’ B compartments in DKO1 have become more active. We also observe that DKO1 have a smaller percentage of the genome organised into topologically-associated domains (TADs) (**Fig. 4d**). Reduced compartment strength and TAD structure suggests a decrease in the number of cells within the population sharing similar chromatin conformation and thus an increase in chromatin conformation heterogeneity. **Fig. 4e** shows representative examples of chromosomes showing reduced integrity of compartmentalisation. Overall, the increase in cell-to-cell heterogeneity of DNA replication is also reflected in increased blurring of 3D compartmentalisation.

**Figure 4:**
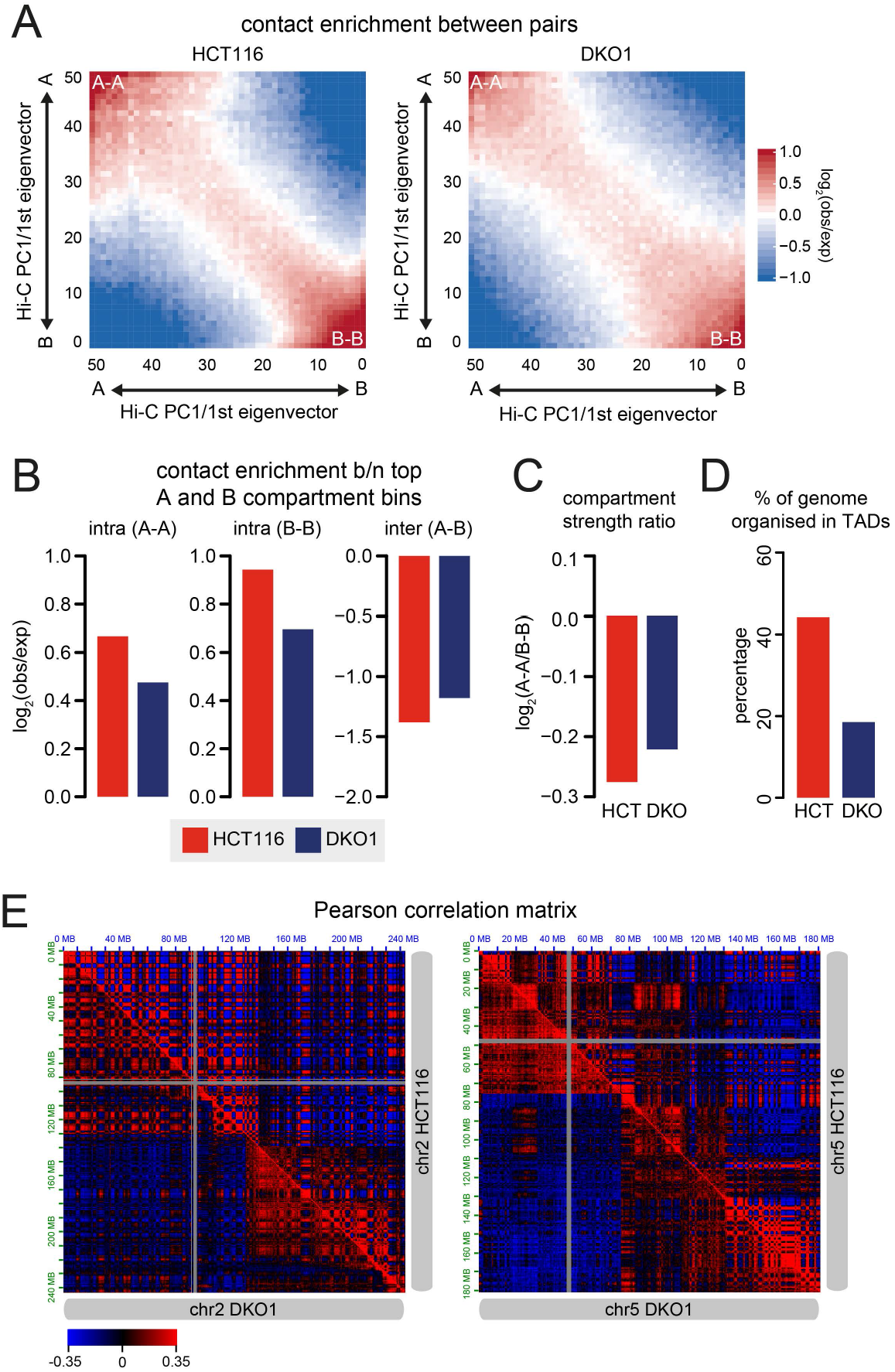
Loss of global methylation reduces integrity of 3D compartmentalisation. **A** Heatmap/Saddle plot showing contact enrichments (log_2_(Obs/Exp)) between pairs of 100kb bins ordered in 50 PC1 quantile groups. **B** Mean contact enrichments within and between A- and B-compartments (top and bottom 20% of PC1 quantiles) are shown. **C** Compartment strength ratio of HCT116 compared to DKO1. Compartment strength ratio is calculated as the log_2_ of the ratio of A-A and B-B mean contact enrichments. **D** Percentage of the genome called as topologically associating domains (TADs). **E** Representative examples of loss of compartmentalisation in DKO1 compared to HCT116. Pearson correlation matrix is shown (Juicebox).

### DNA hypomethylation causes loss of allelic replication

In exploring the decrease in replication timing precision using the weighted average value, we identified a small group of loci that showed a gain of replication timing precision specifically associated with *later* replication timing (**Supp Fig. 5a**). This was contrary to the loss of precision in the rest of the genome. Visual inspection of these loci revealed that they appear to be asynchronously or biphasically replicating loci in HCT116. We therefore asked if DNA hypomethylation can also result in changes in biphasic replication. Biphasically replicating regions occur where the locus is replicated both in early and late timing within the same cell type (Hansen et al., 2010). We called biphasic regions in our HCT116 and DKO1 Repli-Seq datasets using both our weighted variance score and the scoring method from Hansen *et al.* (2010) (**See Methods**). Both methods identified more biphasic regions in HCT116 than DKO1 (**Fig. 5a** and **Supp Fig. 5b**), with ~70-85% of the biphasic regions lost in DKO1 (**Fig. 5b** and **Supp Fig. 5c**). Representative examples of lost biphasic regions are shown in **Fig. 5c** and **Supp Fig. 5d**.

**Figure 5:**
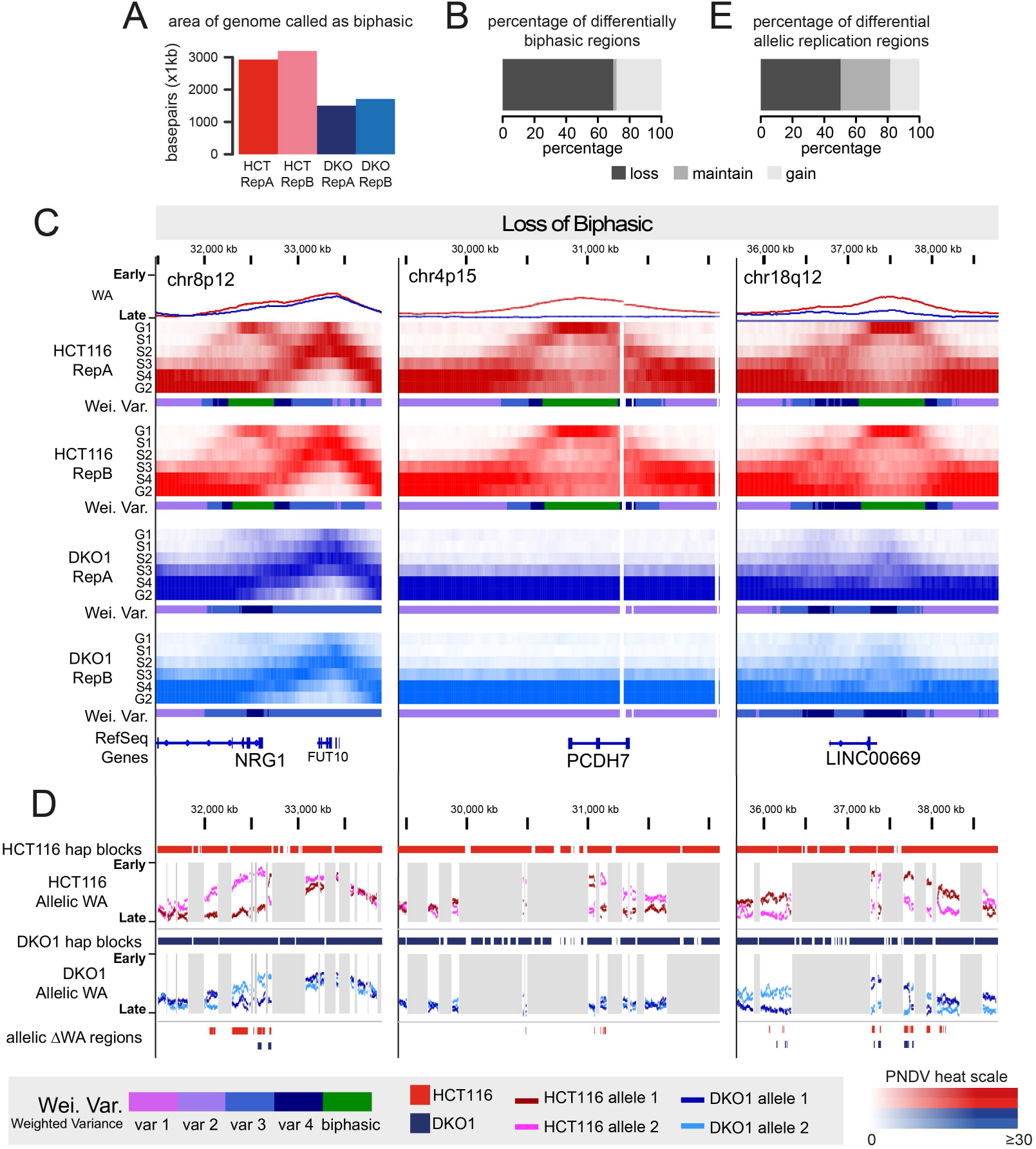
Loss of DNA methylation causes loss of allelic DNA replication. **A** Area of the genome in kilobases (Kb) called as ‘biphasic’ using the weighted variance score. **B** Percentage of biphasic replication regions that are lost, maintained or gained from HCT116 to DKO1 as calculated using the weighted variance score. **C** Examples of regions that changed or maintained biphasic status between HCT116 and DKO1 as calculated using the weighted variance score. All 6 PNDV fractions of each Repli-Seq datasets are shown in the order of G1, S1, S2, S3, S4, and G2 from top to bottom. HCT116 datasets are shown in red and DKO1 datasets are shown in blue. The weighted average replication timing score is shown at the top of each example. Colours indicating bins of the weighted variance score is shown below each set of 6 fractions. Green from the weighted variance score indicate biphasic regions. PNDV value for each fraction is indicated by a heat colour scale. **D** Allelically separated replication timing (WA) scores of the same regions in **C**. Haplotype blocks are shown at the top of each example. Grey shading indicates either no data or a break between haplotype blocks. **E** Percentage of allelically replicating regions that are lost, maintained or gained from HCT116 to DKO1 as calculated using allele-specific replication WA scores.

Biphasic replicating regions in cell population level data may be either due to allelic replication or the presence of two sub-populations with differential replication at the same locus. To determine if biphasic regions show allele-specific replication, we used long-read Nanopore sequencing followed by variant calling and phasing to obtain phased haplotypes for both HCT116 and DKO1 (**See Methods**). Allelically separated replication timing WA scores show that biphasic regions are indeed comprised of one early replicating allele and one late replicating allele in HCT116, and the alleles become more synchronously replicated in DKO1 (**Fig. 5d**, **Supp Fig. 5e,f**). Using allelic replication timing WA scores alone, we confirmed that approximately 50% of allele-specific replicating regions are lost after DNA methylation reduction in DKO1 cells (**Fig. 5e**). Further representative examples of loss of allele-specific replication are shown in **Supp Fig. 6**. The population-level biphasic data together with the Nanopore data show that the hypomethylation predominantly results in loss of asynchronous allele-specific replication.

As imprinting loci are also known to show allelic replication (Kitsberg et al., 1993), we next asked if allele-specific replication in HCT116 occur at imprinted loci. We compared allelically replicated genes against a list of known or predicted imprinted genes (**See Methods**) (Luedi et al., 2007). Of a total of 115 genes located at allelically replicating regions in either HCT116 or DKO1, only 36 were protein coding (**Supp Table 1**) and only two of these, *PRIM2* and *DGCR6*, were found to be in the imprinted gene database. This suggests that the majority of allelic replicating loci in HCT116 are not at developmentally imprinted regions. Interestingly, we found that 16 of the 36 allelically replicating protein-coding genes are also known or predicted to be mono-allelically expressed (Gimelbrant et al., 2007; Savova et al., 2016). Furthermore, many of the allele specific replication genes are reported to be cancer-related (**Supp Table 1**). Notably, three genes that lose allelic replication in DKO1 cells, are also reported to be associated with colon cancer; *NRG1* (Luraghi et al., 2017; Stahler et al., 2017), *PCDH7* (Li et al., 2020; van Roy, 2014) and *DLC1* (Durkin et al., 2007; Peng et al., 2013) (**Fig. 5c,d** and **Supp Fig. 5d,e**). Furthermore, *NRG1* and *PCDH7* are mono-allelically expressed genes and *DLC1* was recently reported to be partially mono-allelically expressed (Gupta et al., 2020). Interesting, we found that loss of allelic replication for these three genes specifically involved loss of the early replicating allele in DKO1. Concordantly, *NRG1*, *PCDH7* and *DLC1* are significantly downregulated in DKO1 compared to HCT116 due to the loss of the early allele (**Supp Fig. 5g**). This suggests that DNA hypomethylation promotes an alteration of allelic replication timing and reduced expression of some key cancer-related genes.

### Chromatin modifications and gene expression changes following DNA hypomethylation are associated with altered 3D genome architecture

Chromatin changes in repressive heterochromatin histone marks (i.e. H3K27me3, H3K9me3) have been shown to occur following loss of DNMT expression (Espada et al., 2004; Reddington et al., 2013; Saksouk et al., 2014). Furthermore, heterochromatin has been shown to be important in establishing and maintaining global nuclear organisation and compartmentalisation (Belaghzal et al., 2019; Falk et al., 2019; MacPherson et al., 2018). Therefore, we asked where co-ordinated histone modification changes occur in the genome after DNA hypomethylation in DKO1 cells, and if this relates to higher order genome architecture. To do this, we called chromHMM states for both HCT116 and DKO1 and found that the majority of state changes between HCT116 and DKO1 were the result of a gain of H3K4me3, H3K4me1 and H3K27me3, and a loss of H3K36me3 and H3K9me3 (**Supp Fig. 7a,b, See Methods**). These changes can also be observed in the histone marks themselves (**Fig. 6a**). We observed an overall increase in H3K27me3 and H3K4me3, and a loss of H3K9me3 mostly at late-replicating loci. Surprisingly, gain of H3K4me3 occurs at both early- and late-replicating loci. The gain of H3K27me3 and loss of H3K9me3 reflects our previous observations at hypomethylated late-replicating regions in prostate and breast cancer cells (Du et al., 2019). Moreover, the loss of heterochromatin mark H3K9me3 in late replicating regions agrees with the loss of chromatin compartmentalisation integrity (**Fig. 4**).

**Figure 6:**
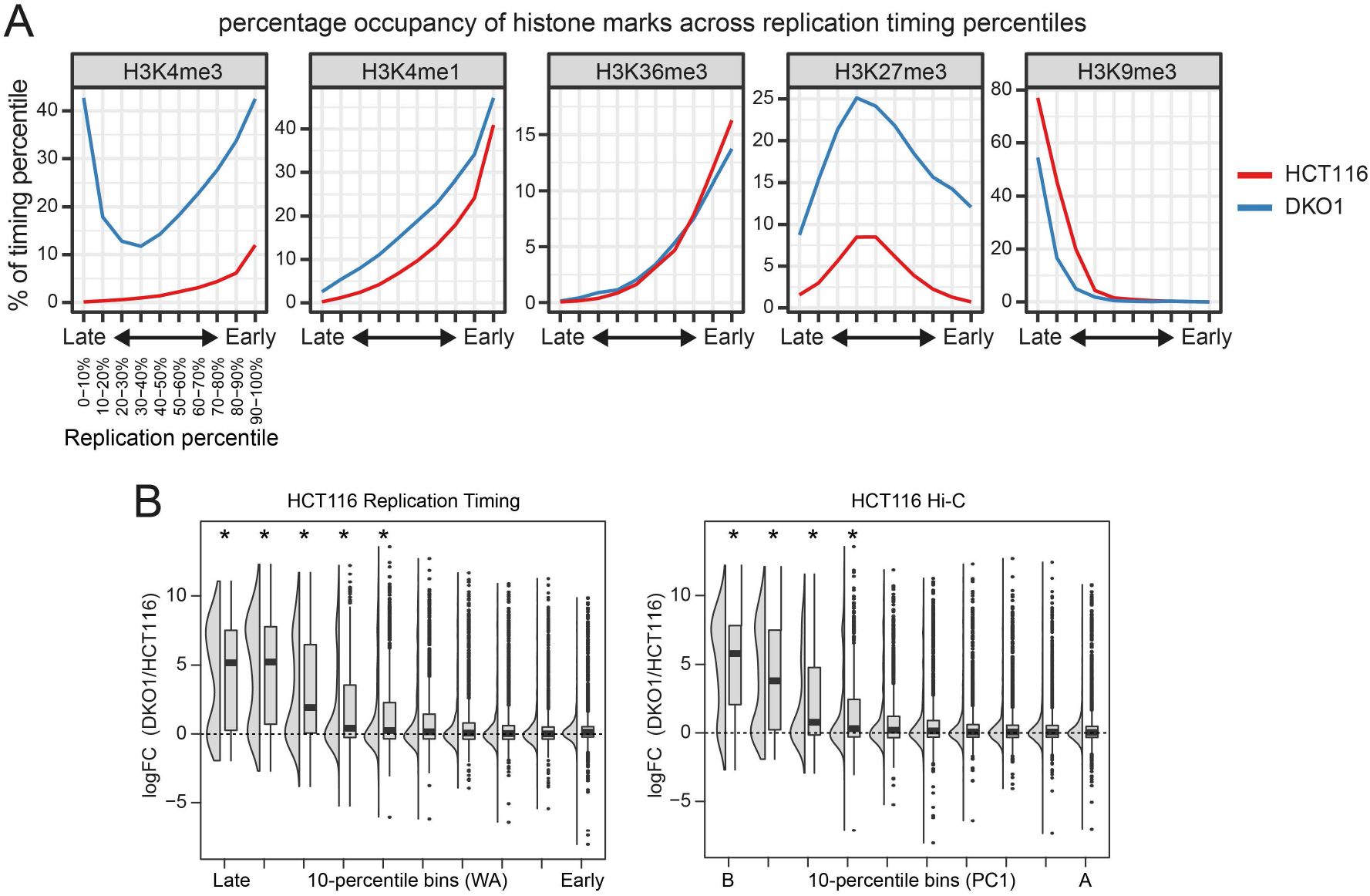
DNA replication timing stratifies chromatin changes between HCT116 and DKO1. **A** Percentage occupancy of histone marks for 1kb loci across replication timing bins in HCT116 and DKO1. HCT116 data is shown in red and DKO1 data is shown in blue. Replicated peaks were used where replicates were available (H3K4me3, H3K4me1, H3K27me3, H3K9me3). **B** Boxplots and density plots of differential expression (logFC) of genes within each of 10-bins of replication timing (WA) or Hi-C PC1 values of HCT116. Asterisks indicate significance in one-tailed Mann-Whitney-Wilcoxon test of each bin against genome-wide differential expression, where the alternative ‘greater’.

Next, to explore whether gene deregulation, between HCT116 and DKO1, is related to replication timing and chromatin conformation changes we examined differential gene expression. We found 2291 upregulated and 150 downregulated genes in DKO1 compared to HCT116 (**Supp Fig. 7c**). Overall, genes that replicated *earlier* and located in B-A shifted compartments (positive ΔPC1) are associated with upregulation and genes that replicated *later* and located in A-B shifted compartments (negative ΔPC1) are associated with downregulation of expression (**Supp Fig. 7d**). Furthermore, we found that genes located within late replication timing and B-compartments (negative PC1 values) show the most expression upregulation in DKO1 (**Fig. 6b, Supp Fig. 7e**). These results suggest that loss of DNA methylation is leading to aberrant gene activation in late-replicating B-compartments.

### Gain of broad H3K4me3 in H3K9me3 domains after hypomethylation may protect against genome reorganisation

One of the most significant changes in chromHMM states between HCT116 and DKO1 is the change from H3K9me3 enrichment (heterochromatin, Het) to H3K4me3 enrichment (active TSS, TssA) chromatin state. This was notable and appears to occur in large domains that coincide with late-replicating domains in both HCT116 and DKO1 (**Fig. 7a**, **Supp Fig. 8a**). These regions have maintained H3K9me3 between HCT116 and DKO1. However, DKO1 also displays a low but broad gain in H3K4me3 enrichment across the same region, thereby contributing to the change in chromHMM state from Heterochromatin to TssA (**Fig. 7a**, **Supp Fig. 8a**). The broad H3K4me3 domains do not resemble the typically observed punctate H3K4me3 peaks and interestingly, the broad enrichment is only observed when H3K9me3 is maintained from HCT116 to DKO1 (**Fig. 7a**, **Supp Fig. 8a**, beige shading), but not when H3K9me3 is lost in DKO1 (**Fig. 7a**, **Supp Fig. 8a**, blue shading).

**Figure 7:**
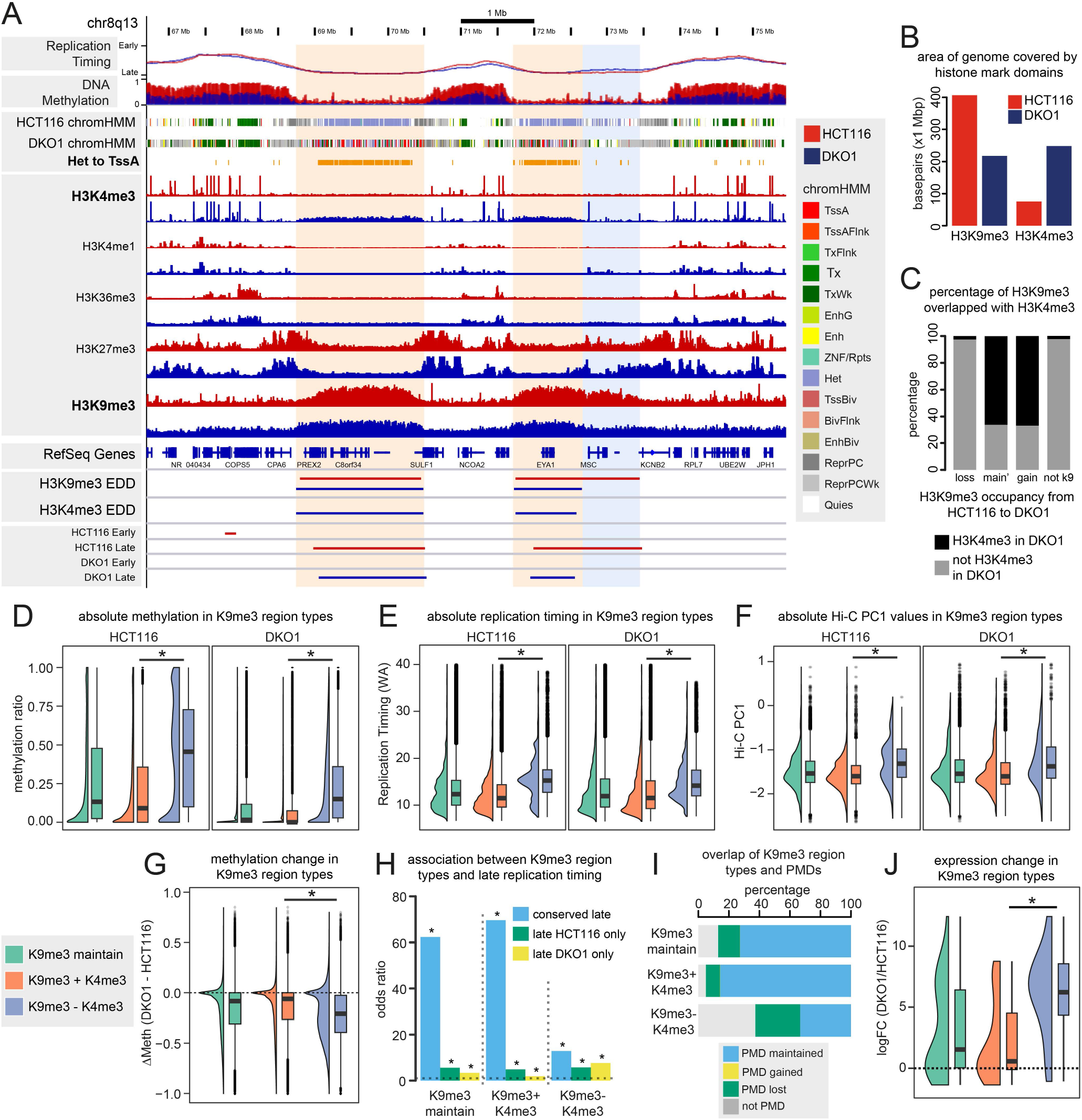
Non-canonical H3K9me3 and H3K4me3 bivalent domains occur at conserved late-replicating B-compartments and maintains expression silencing. **A** Representative example of broad H3K9me3 and H3K4me3 enrichment. All HCT116 datasets are shown in red and all DKO1 datasets are shown in blue. The ‘H09_Het to D01_TssA’ track shows loci of the chromatin state transition (See **Supp Fig. 7**). Beige shading indicates H3K9me3 domains maintained from HCT116 to DKO1. Blue shading indicates H3K9me3 domains lost from HCT116 to DKO1. **B** Area of the genome in Megabases (Mb) covered by H3K9me3 and H3K4me3 board domains. **C**Percentage occupancy of H3K9me3 and H3K4me3 broad histone domains between HCT116 and DKO1. **D,E,F** Absolute DNA methylation levels, absolute replication timing levels and absolute Hi-C PC1 levels in H3K9me3/H3K4me3 region types. For **D**, **E** and **F**, asterisks indicate that the K9me3+K4me3 regions are less than the K9me3-K4me3 regions (*p* < 0.05, one-tailed Mann-Whitney-Wilcoxon tests). **G** DNA methylation change of H3K9me3/H3K4me3 region types. Horizontal dotted lines indicates no change (ΔMeth = 0). Asterisks indicate that the K9me3+K4me3 region is greater than the other categories (*p* < 0.05, one-tailed Mann-Whitney-Wilcoxon tests). **H** Association between H3K9me3/H3K4me3 region types and either conserved late domains, HCT116-only late domains or DKO1-only late domains. Only significant associations are shown (Fisher’s exact test, FDR < 0.05). **I** Overlap of H3K9me3/H3K4me3 region types with maintained, lost and gained PMDs between HCT116 and DKO1. **J** Differential expression (logFC, DKO1/HCT116) of transcripts within H3K9me3/H3K4me3 region types based on promoter overlaps. Horizontal dotted line indicates no change (logFC = 0). Asterisks indicate that the K9me3+K4me3 is less than K9me3-K4me3 (*p* < 0.05, one-tailed Mann-Whitney-Wilcoxon tests).

To examine the H3K9me3/H3K4me3 patterns genome-wide, we called broad domains of H3K9me3 using Enriched Domain Detector (EDD) (Lund et al., 2014). Comparing maintained, lost or gained H3K9me3 regions between HCT116 and DKO1, we found that maintained H3K9me3 regions showed the highest increase in H3K4me3 enrichment in DKO1 (**Supp Fig. 8b**). There is no obvious increase of H3K4me1, H3K36me3 or H3K27me3 in the H3K9me3 maintained regions (**Supp Fig. 8c**) suggesting this enrichment is H3K4me3-specific.

We next called broad domains of H3K4me3 and tabulated the co-occupancy of H3K9me3 and H3K4me3 between HCT116 and DKO1 (**See Methods**). These broad domains can be seen in **Fig. 7a**. Compared to HCT116, DKO1 has lost half of the broad H3K9me3 domains but has more than doubled the area of the genome called as a broad H3K4me3 domain (**Fig. 7b**). Approximately 66% of H3K9me3 domains in DKO1 (maintained or gained compared to HCT116) overlap with newly acquired H3K4me3 domains in DKO1 (**Fig. 7c**). In contrast, there were minimal novel H3K4me3 domains in regions that lacked H3K9me3 domains (**Fig. 7c**). Altogether, these results show that after DNA methylation loss, regions of the genome that maintain H3K9me3 become broadly marked by H3K4me3 to form non-canonical bivalent domains.

To further characterise these bivalent domains (H3K9me3+H3K4me3), the H3K9me3 maintained domains were further separated into those with or without H3K4me3 in DKO1 (‘K9me3+K4me3’ vs. ‘K9me3-K4me3’). H3K9me3 is highly abundant in late replication timing loci in cancer cells (Du et al., 2019), therefore regions with H3K9me3 are expected to have low DNA methylation and low Hi-C PC1 values. However, we observed that H3K9me3+H3K4me3 domains are consistently less methylated, display later timing and have lower Hi-C PC1 values than H3K9me3-H3K4me3 domains (**Fig. 7d-f**). We also observed reduced methylation loss (**Fig. 7g**), which appears to be due to the non-canonical bivalent H3K9me3+H3K4me3 domains having very low initial methylation in HCT116 compared to H3K9me3-H3K4me3 domains (**Fig. 7d**), and essentially becoming completely unmethylated in DKO1. The gain of H3K4me3 may therefore be a response to a complete methylation loss at these specific regions in DKO1 cells.

Non-canonical bivalent H3K9me3+H3K4me3 domains also show more association with conserved late replication between HCT116 and DKO1, compared to H3K9me3-H3K4me3 (**Fig. 7h**). Agreeing with the overlap with conserved late regions, H3K9me3+H3K4me3 domains also overlap more with maintained PMDs compared to H3K9me3-H3K4me3 domains (**Fig. 7i**). This suggests that in response to global DNA methylation loss, late replication and genome organisation is maintained by the formation of these non-canonical broad bivalent H3K9me3+H3K4me3 domains.

The increase in H3K4me3 may suggest a gain of transcriptional activity within H3K9me3 maintained regions. Therefore, we examined the transcriptional activity of genes within H3K9me3 domains with and without H3K4me3. Interestingly, bivalent H3K9me3+H3K4me3 domains showed lower expression upregulation compared to H3K9me3-H3K4me3 domains (**Fig. 7j**). Rather than the expected gain of transcriptional activity, these results suggest that the H3K9me3+H3K4me3 co-occupancy protects these regions against an overall gene activation that occurs upon global hypomethylation in DKO1 cells (see **Supp Fig. 7c**). Altogether, these results suggest that after DNA methylation loss, gain of broad H3K4me3 occupancy in H3K9me3-maintened domains, may potentiate maintenance of late replication timing and PMD structure, and protect against aberrant gene activation.

## Discussion

DNA hypomethylation is a hallmark of cancer cells but its impact on 3D genome architecture is not well understood. Therefore, we were interested to determine the direct effect of DNA hypomethylation on the extent of DNA replication timing and 3D genome organisational changes. Using a colorectal cancer cell line model of DNA hypomethylation, we show that loss of DNA methylation results in a genome-wide increase in replication timing heterogeneity and a corresponding loss of 3D genome compartmentalisation. The genome-wide loss of higher order chromatin stability was accompanied by *earlier* replication and a shift in nuclear organisation towards active A-compartments at the boundaries of partially methylated domains (PMDs). We further identified that DNA hypomethylation causes a notable loss of allelically replicating regions across the genome. Lastly, we found that DNA hypomethylation was associated with gain of non-canonical bivalent domains comprised of H3K4me3 and H3K9me3, which we suggest may provide protection against loss of late replicating B-compartment PMDs and aberrant gene activation. Altogether, this study has revealed that the methylome is critical for the maintenance of higher order genome architecture.

There have been limited studies to date on the consequences of disrupting the methylome on 3D genome organisation. Imaging studies have reported de-condensation at chromocenters after *Dnmt1* KO in mouse embryonic fibroblasts (Casas-Delucchi et al., 2012) and an increase in DNase-I sensitivity of the inactive X chromosome after 5-azacytidine treatment of gerbil lung fibroblasts (Jablonka et al., 1985). More recently, loss of compartmentalisation was found in senescent fibroblasts with *DNMT1* knockdown (Sati et al., 2020). In contrast, a prior study using the HCT116 DKO1 hypomethylation cell model did not find an increase in genome-wide open chromatin, suggesting no chromosomal de-condensation (Pandiyan et al., 2013). However, our findings of global de-compartmentalisation after hypomethylation further support that DNA methylation is required to maintain the stability of 3D genome organisation.

Importantly, no studies thus far have examined the global consequence of hypomethylation on DNA replication timing. Using single cell sequencing, we show that large scale cell-to-cell heterogeneity occurs in replication timing in the HCT116 DKO1 cell model, suggesting that DNA hypomethylation has caused an ‘erosion’ of the precise regulation of replication timing. We speculate two related mechanisms that may drive the increased heterogeneity in replication timing. First, it is possible that a loss of DNA methylation affects the regulation of replication origins leading to loss of replication timing precision. Studies have previously reported mammalian replication origins to be located at or near CpG-islands (Cadoret et al., 2008; Cayrou et al., 2011; Sequeira-Mendes et al., 2009) and that DNA methylation can stabilise interactions between H3K9me3 and origin recognition complex (ORC) proteins (Bartke et al., 2010). In DNA methyltransferase KO studies in mouse and human cells, loss of DNA methylation results in loss of H3K9me3 (Espada et al., 2004; Saksouk et al., 2014). However, the targeted removal of H3K9me3 by its demethylase, Kdm4d, at replication origins is essential for origin initiation and elongation (Wu et al., 2017). Therefore, DNA methylation may play a role in stabilising origin location by preventing ectopic origin activation, and hence loss of methylation would lead to replication timing imprecision via disorganised origin activation. Further studies will be needed however to finely elucidate a role of DNA methylation in the control of replication origins. Secondly, our findings that late-replicating B-compartment regions show the most increase in expression, as well as loss of the heterochromatic mark H3K9me3, support that the late-replicating heterochromatic regions are compromised after DNA methylation loss. Heterochromatin, particularly H3K9me3 and its reader HP1, have been shown to be important in establishing and maintaining global chromatin conformation, phase separation and compartmentalisation (Falk et al., 2019; Larson et al., 2017; MacPherson et al., 2018). Indeed, loss of the H3K9me3 methyltransferases (*Suv39h1,2*) or HP1 also lead to *earlier* replication timing of chromocenters and centromeric repeats (Schwaiger et al., 2010; Wu et al., 2006), similar to loci-specific studies of DNA methylation loss. Therefore, loss of H3K9me3 in DKO1 may destabilise the organisation of the genome, hence, driving the global de-compartmentalisation and decreased replication timing stability we observe.

We further identified specific genomic loci, covering ~3-10% of the genome, that showed a complete switch in replication timing and chromatin conformation after DNA methylation loss. The majority of 3D genome architectural changes we identified involved switching to *earlier* DNA replication timing at the boundaries of partially methylated domains (PMDs), accompanied by a similar shift towards active A-compartment values in the Hi-C PC1 score (**Fig. 2g**). Loss of DNA methylation has been reported to result in *earlier* onset of DNA replication at a number of heterochromatic loci such as chromocenters (Casas-Delucchi et al., 2012), pericentromeric major satellite repeats (Jorgensen et al., 2007) and the inactive X chromosome (Hansen et al., 2000; Jablonka et al., 1985). PMDs are known to be marked by H3K9me3/H3K27me3 heterochromatin domains (Hon et al., 2012; Salhab et al., 2018), thus agreeing with prior loci-specific studies. However, our results show that *earlier* replication resulting from loss of DNA methylation is more common than previously described and occurs throughout the genome. We further observed that loss of methylation at PMD boundaries results in a ‘blurring’ of the methylation boundaries and shrinking of the PMD. Further exploration of gene expression, histone modification, gene structure or transcription factor occupancy differences between maintained and changed PMD boundaries may reveal the exact mechanisms of why only some PMDs are affected.

We next found that discrete loci showing allele-specific replication in HCT116 are lost after DNA hypomethylation in DKO1. Early studies reported that DNA methylation imprinted regions can show allelic-specific replication, for example at imprinting loci located on X chromosome (Takagi and Oshimura, 1973), *IGF2*, *H19* and *SNRPN* (Kitsberg et al., 1993). DNA hypomethylation and associated loss of imprinting commonly occurs in cancer (Jelinic and Shaw, 2007; Robertson, 2005). Therefore, the observation of allele-specific replication in the colorectal cancer cells, HCT116, was unexpected. However, similar to other studies (Hansen et al., 2010; Rivera-Mulia et al., 2018), we found that few allelically replicating loci contain known or predicted imprinted genes, suggesting that allelic replication may be a separate regulatory mechanism to classic gene imprinting. Interestingly, a substantial proportion of the allelically replicating regions contain genes previously identified as showing monoallelic expression (Gimelbrant et al., 2007). Since the majority of allele-specific replicating loci in HCT116 lost their allelic replication after DNA hypomethylation, we suggest that DNA methylation also plays a role in the regulation allele-specific replicating loci that are not imprinted loci. In agreement, loss of allelic replication at key cancer-related genes (*TP53* and *RB1*) was found in cancer patient lymphocytes treated with the demethylation agent 5-azacytidine (Dotan et al., 2004, 2008; Nagler et al., 2010). There is also evidence for DNA methylation involvement in non-imprinted monoallelic expression in both humans (da Rocha and Gendrel, 2019; Schalkwyk et al., 2010) and mice (Gupta et al., 2020). We further identified that loss of allelic replication was associated with expression down-regulation at three colon cancer-related gene loci that are also monoallelically expressed. Monoallelic expression has also been reported to differ in frequency between grades of brain tumours (Walker et al., 2012) and occur at different genes in colorectal tumours compared to normal tissue (Liu et al., 2018), suggesting that cancer cells can acquire ectopic allele-specific expression during transformation. Together, our results suggest that altered allelic replication, induced by global changes in DNA methylation, may be a mechanism of gene deregulation within cancer cells.

It is well established that bivalent H3K4me3-H3K27me3 states commonly occur in normal cells at silent CpG island promoters, and that in cancer, bivalency is lost when these promoters gain DNA methylation (Baylin and Jones, 2016). We were therefore intrigued to find that global loss of DNA methylation in DKO1 cells was associated with the formation of non-canonical bivalent domains of broad H3K4me3 enrichment in H3K9me3 domains genome-wide. Broad H3K4me3 ‘mesas’ have previously been described in senescent cells and form over LADs (Shah et al., 2013), and examples of broad H3K4me3/H3K9me3 loci have previously been noted in DKO1 cells (Lay et al., 2015). However, here we further demonstrated that these H3K9me3+H3K4me3 domains specifically maintain late replication and PMD structure between HCT116 and DKO1, and show the least aberrant gene activation, despite gain of the H3K4me3 active mark. More recently, broad H3K4me3 domains have also been identified in mouse oocytes (Dahl et al., 2016; Liu et al., 2016; Zhang et al., 2016). These non-canonical H3K4me3 domains appear following the wave of DNA methylation erasure that occurs during reprogramming of primordial germ cells, and form during a period of genomic silencing in late stage (MII) mature oocytes (Dahl et al., 2016; Zhang et al., 2016). Similar to our findings, these non-canonical H3K4me3 domains also co-occur with PMDs (Zhang et al., 2016) and are anti-correlated with DNA methylation (Dahl et al., 2016). Active *de novo* methylation by DNMT3A and -3B is required to protect regions *against* acquiring this form of H3K4me3 in oocytes (Hanna et al., 2018). Therefore, it is possible that the knockout of *de novo* methyltransferase *DNMT3B* may be the main driver of H3K4me3 domain formation in DKO1 cells. The H3K4me3 domains further function in oocytes to maintain zygotic genome silencing (Zhang et al., 2016), suggesting that the ectopic H3K9me3+H3K4me3 bivalent domains we observe in DKO1 cells may also be protecting late-replicating H3K9me3-marked domains from spurious activation in response to extreme DNA methylation loss. Altogether, this data suggests that KO of *DNMT1* and *DNMT3B* in DKO1 cells may be creating a similar demethylation event that occurs in late stage (MII) oocytes, subsequently causing the acquisition of non-canonical H3K4me3 domains.

In summary, our results demonstrate that DNA hypomethylation, a hallmark of cancer cells, leads to disruption of DNA replication timing precision, cell-to-cell heterogeneity and higher-order genome reorganisation. The resulting 3D genome heterogeneity may create further opportunities for clonal selection during tumourigenesis. It will be important in future studies to use more single cell epigenomic approaches to understand the temporal relationship between changes in the epigenome and the 3D genome architecture during tumour progression.

## Acknowledgements

We thank members of the Clark Laboratory for helpful discussions and careful reading of the manuscript. We thank Prof. Stephen Baylin for kindly providing the HCT116 and DKO1 cells. We thank Phillippa C. Taberlay and Yi Cai for advice with tissue culture. S.J.C. is a National Health and Medical Research Council (NHMRC) Senior Principal Research Fellow #1063559. Q.D. is a NHMRC Investigator Grant recipient #1177792. This work was supported by NHMRC Project Grants APP1147974; APP1163939 (S.J.C) and was partially enabled by a Cancer Institute New South Wales (CINSW) equipment grant REG181268 (M.A.S.). The contents of the published material are solely the responsibility of the administering institution and individual authors and do not reflect the views of the NHMRC.

## Author Contributions

Conception and Design, Q.D. and S.J.C.; Methodology, Q.D., M.B., C.E.C., D.K., and J.E.P.; Investigation, Q.D., G.C.S., E.C., S.S.N., D.K., K.B., and J.A-K.; Analysis, Q.D., P.L.L., J.M.F., N.J.A., C.M.G., E.Z., M.B. and J.A-K.; Interpretation of Data, Q.D., K.S., J.A-K. and S.J.C.; Writing – Original Draft, Q.D. and S.J.C.; Writing – Review & Editing, Q.D., C.S., J.A-K. and S.J.C.; Resources, M.A.S., J.E.P., and I.A.D.; Funding Acquisition, S.J.C. and C.S.

## Methods

### HCT116 and DKO1 cell model

HCT116 and DKO1 cells were kindly provided by Prof. Stephen Baylin (The Johns Hopkins School of Medicine, The Sidney Kimmel Comprehensive Cancer Center). HCT116 human colorectal cancer cells were cultured in McCoy’s 5A modified medium (Gibco, #16600-082) supplemented with heat inactivated foetal bovine serum (10%, Gibco, 16000-044) at 37°C and 5% CO_2_. Cells at 70-80% confluency were rinsed in PBS (1x Phosphate-buffered saline, Gibco, #14190-144) and trypsinised in Trypsin-EDTA (0.05%, Gibco, #15400054). Trypsin was inactivated using equal volume growth medium and cells were pelleted at 250xg for 5 min. Cells were resuspended in growth medium and typically split 1:8. HCT116 with double knockouts (KO) in DNMT1 and DNMT3B (DKO1) (Rhee et al., 2002) were selected in growth medium supplemented with hygromycin (0.05 mg/mL, Gibco, #10687-010) and geneticin (0.1 mg/mL, Gibco, #10131) for 1 week after thawing to ensure a pure KO population, then further cultured without selection. DKO1 cells were subcultured following the same protocol as for HCT116 above, but typically at a 1:4 split. HCT116 and DKO1 cells were validated for double knock out of *DNMT1* and *DNMT3B* using western blot and expression qRT-PCR (**Supp Fig. 9**).

### Expression qRT-qPCR of *DNMT* genes

RNA was extracted from cultured cells using TRIzol (Life Technologies, #15596018) and DNaseI treated (NEB, #M0303) as per manufacturer’s instructions. cDNA was generated from 500 ng of RNA using SuperScript III reverse transcriptase (Invitrogen, #18080-044) and random hexamers according to manufacturer’s instructions. 1 μL of RNA sequins (Hardwick et al., 2016) (mix A, 1:100 dilution) was spiked in to RNA samples prior to making cDNA to be used as negative controls for qRT-PCR. Primers to *DNMT1* were designed to exons present in both the full transcript and the hypomorphic product present in DKO1 cells (**Supp Table 2**). Primers to *DNMT3A* and *-3B* were designed to capture the majority of transcripts annotated by the GENCODE genes v19 track in UCSC Genome Browser (Kent et al., 2002) (**Supp Table 2**). All primer pairs were designed over an intron to avoid genomic products.

### Western blotting for DNMT proteins

Whole cell protein lysates were prepared by resuspending scaped cells in modified RIPA buffer (50 mM Tris-HCL pH 7.5, 150 mM NaCl, 1 mM EDTA, 1 mM NaF, 1 mM Na_3_VO_4_, 1% Igepal, 0.25% Sodium-deoxycholate) with protease inhibitors and incubated on ice for 30 min with vortexing every 5 min. Lysate was sonicated for 15s at 25% amplitude with a microtip using the QSonica (Q55) and stored at −80°C. Protein concentration was determined using a BCA assay (Pierce, #23225) according to manufacturer’s instructions. Protein lysate was prepared using NuPAGE® LDS sample buffer (Life Technologies, NP0007) and NuPAGE® sample reducing agent (Life Technologies, NP0004) followed by heating at 95°C for 10 min. Samples were resolved by gel electrophoresis using the NuPage Bis-Tris 4-12% precast gel system according to the manufacturer’s instructions (Life Technologies). Western blot transfer was carried out according to the manufacturer’s instructions with the SureLock X-cell system (ThermoFisher Scientific) using transfer buffer containing 20% methanol content. Antibodies used are as follows: N-terminal DNMT1 – Sigma-Aldrich #D4692; C-terminal DNMT1 – Abcam #ab92314; DNMT3A – Active Motif ##39206; DNMT3B – Active Motif #39207; GAPDH – Thermo Fisher #AM4300. Western blots (WBs) were then treated with Western Lightning Plus-ECL (Perkin Elmer, #NEL103E001) before developing on Super Rx Fuji Medical X-Ray Film (Fujifilm, #4741019236) using the Konica Tabletop X-Ray Film Processor (#SRX101A). Developed film was scanned using the Epson Perfection V800/850 scanner. ImageJ was used to quantitate WB films following the *Gel Analysis* method outlined in their documentation. Band densities of the protein of interest were divided by the loading control and converted to a percentage of the highest ratio (most dense band compared to loading control). Average and standard error of the mean were calculated for replicates.

### Whole genome bisulphite sequencing and processing

200 ng of DNA was bisulphite converted using the EZ DNA Methylation-Gold Kit (Zymo, #D5005) according to manufacturer’s instructions. Input DNA was spiked with unmethylated lambda DNA (0.5%) (Promega, #D1521). Replicate bisulphite libraries were generated with the CEGX TrueMethyl® Whole Genome Kit (CEGX, #CEGXTMWG, v3.1) according to manufacturer’s instructions. Libraries were sequenced on the Illumina X Ten. Sequencing reads from WGBS data were aligned to the human genome using v1.2 of an internally developed pipeline Meth10X (Nair et al., 2018). This is publicly available and can be downloaded from https://github.com/luuloi/Meth10X. The pipeline backbone is built based on workflow control Bpipe (v0.9.9.2) (Sadedin et al., 2012). Briefly, adaptor sequences were removed using in-house bash script in paired-end mode following library prep kit guide. Bwa-meth (v0.20) (Pedersen et al., 2014) was then used to align reads to hg19 using default parameters. The generated bam files were marked for duplicates using Picard (v2.3.0) (http://broadinstitute.github.io/picard). Bam files were then quality checked using Qualimap (v2.2.1) (Okonechnikov et al., 2016). Count tables of the number of methylated and unmethylated bases sequenced at each CpG site in the genome were constructed using MethylDackel (https://github.com/dpryan79/MethylDackel) and Biscuit (https://github.com/zwdzwd/biscuit). Biscuit was used to call SNPs that were discounted from the final table. Sequencing metrics can be found in **Supp Table 3**. Partially methylated domains (PMDs) were called using MethPipe (v3.4.2) (Song et al., 2013).

### Repli-Seq data generation and processing

Repli-Seq was performed in duplicate for each cell line as previously described with slight modifications (Du et al., 2019). Briefly, cells were labelled with BrdU (50 μM, Sigma, #B5002) for two hours. Labelled cells were sorted into 6 fractions across the cell cycle (G1b, S1, S2, S3, S4, G2M) as per protocol on the FACS Aria III. DNA extraction and BrdU-labelled DNA immunoprecipitation were performed with anti-BrdU antibody (40 μL of 25 μg mL^−1^, BD Pharmingen, #555627). Validation of BrdU immunoprecipitation was carried out using qRT-PCR on known Early (*BMP1*) and Late (*DPPA2*) loci, and compared to a fractionation negative control (MITO) (**Supp Fig. 1a,b**). As the mitochondrial genome (MITO) replicates independently of the nuclear genome, it should not show differences between S-phase fractions. 10 ng of ssDNA was used as input for the Epicenter EpiGnome™ Methyl-Seq Kit (Illumina, EGMK81312, now called the TruSeq® DNA Methylation kit) and processed according to manufacturer’s instructions. The ssDNA was not bisulphite converted prior to library preparation. Libraries were sequenced on the HiSeq 2500 as 50bp single-end reads. Full sequencing outputs can be found in **Supp Table 4**.

Replication timing weighted average (WA) scores were calculated as previously described (Du et al., 2019). WA values for replicates of HCT116 and DKO1 were highly correlated (r^2^ values >0.99) (**Supp Fig. 1c**). The distributions of WA scores were comparable to the WA distributions in other normal and cancer cell Repli-Seq datasets (n=16) (**Supp Fig. 1d**). To get a single score per cell line, replicate HCT116 and DKO1 weighted average (WA) replication timing scores were quantile normalised and the replicates averaged. Early- and late-replicating regions were defined as those regions in the top and bottom 10% of WA scores in both cell lines. This definition gives upper and lower limits of 77.76 and 16.07 respectively (i.e. early regions have WA > 77.76 and late regions WA < 16.07). The limits were rounded to 78 and 16 for downstream analyses. WA thresholds for a change in timing were were calculated as previously described (Du et al., 2019). Differences in WA that are larger than +/− 15 ΔWA are therefore considered to show a robust change in replication timing (**Supp Fig. 1e,f**). To identify domains of loci with changed replication timing, we merged all loci within 50kb that had |ΔWA| > 15. This approach gave 176 *earlier* domains and 117 *later* domains called between HCT116 and DKO cells.

### Defining partially methylated domain boundary shifts

Partially methylated domains (PMDs) were called from WGBS data using the package MethPipe (v3.4.2) (Song et al., 2013). To call PMD regions for HCT116 and DKO1, first, replicate PMD regions were merged and any regions smaller than 50kb were removed. Then regions were merged if they were within 50kb of each other. The cut-off of 50kb was used because this is the resolution of the replication timing data. To define PMD boundary shifts, we first calculated which PMD regions overlap between HCT116 and DKO1, then calculated the distance between the 5’ start coordinate of the HCT116 PMD to the 5’ start coordinate of the DKO1 PMD. The same was repeated for the 3’ end coordinate. The 5’ and 3’ coordinates were then categorised by the degree of shift inwards from the HCT116 to DKO1 into 4 categories, <50kb, 50-200kb, 200-500kb, >500kb. Due to the low numbers of regions within the larger shift categories, 5’ and 3’ regions were pooled prior to plotting the changes in replication timing and DNA methylation.

### Hi-C library preparation

Hi-C data was generated using the Arima-HiC kit, according to the manufacturers protocols (Cat. #A510008). Briefly, cells were cross-linked with 2% formaldehyde to obtain 1-5 μg of DNA per Hi-C reaction. The Arima kit uses two restriction enzymes: ^GATC (DpnII), G ^ANTC (N can be either of the 4 genomic bases) (HinfI), which after ligation of DNA ends generates 4 possible ligation junctions in the chimeric reads: GATC-GATC, GANT-GATC, GANT-ANTC, GATC-ANTC. Hi-C libraries were prepared using the KAPA Hyper Library Prep Kit with a modified protocol provided by Arima with 12 PCR cycles for library amplification as required. Hi-C libraries were sequenced on Illumina HiSeq X Ten in 150bp paired-end mode.

### Hi-C data processing

HiC-Pro (Servant et al., 2015) (v2.11.4) was used to align and filter the Hi-C data, identify chromatin interactions, and generate Hi-C heatmaps. To generate filtered Hi-C contact matrices, the Hi-C reads were aligned against the human reference genome (hg19) and corrected using the ICE “correction” algorithm (Imakaev et al., 2012) built into HiC-Pro. Statistics on the number of read pairs, valid interactions and interactions in cis are presented in **Supp. Table 5**.

Contact matrices used in down-stream analysis were Knight-Ruiz (KR)-normalized using JuiceBox tools (Durand et al., 2016a; Durand et al., 2016b) using Hi-C contact matrices in .hic format generated by hicpro2juciebox script in HiC-Pro as input. Obtained Hi-C matrices and Pearson correlation matrices were visualised in JuiceBox (Durand et al., 2016a).

### TADs

KR-normalized contact matrices were retrieved from Juicer for all chromosomes at 40Kb resolution and TADs were identified using TADtool with the insulation score algorithm (Kruse et al., 2016). We called TADs with a window size value of 103kb and a TAD cutoff of 30. We found that these parameters show good agreement between identified TADs and visual inspection of Hi-C datasets in JuiceBox and TADtool.

### A/B Compartments

Compartment analysis was performed using the Homer pipeline (Heinz et al., 2010) (Heinz et al., 2010) (v4.6) with Hi-C KR-normalised contact matrices as input. Homer performs a principal component analysis of the normalized interaction matrices and uses the PC1 values to predict regions of active (A-compartments) and inactive chromatin (B-compartments). Homer works under the assumption that gene-rich regions have similar PC1 values, while gene-poor regions show differing PC1 values and assigns compartment status based on genome-wide gene density. To identify compartments that switched between HCT116 and DKO1, we getHiCcorrDiff.pl pipeline was used to correlate the interaction profile of each locus in the HCT116 to the interaction profile of that same locus in the DKO1.

### Defining domains of nuclear reorganisation

We initially tried to define nuclear reorganisation through A/B compartment switching using compartments defined by HOMER. However, we observed that a large proportion of compartment switches were centred around PC1 values of zero, close to the A/B boundary; 47.90% of A to B switches occur where HCT116 PC1 < 0.5 and DKO1 PC1 > −0.5, and 43.24% of B to A switches occur where HCT116 PC1 > −0.5 and DKO1 PC1 < 0.5 (**Supp Fig. 2e**). This indicates that although compartment switches do cross the midline that defines A vs. B compartments, a large proportion of these switches show little difference in PC1 values between HCT116 and DKO1 (**Supp Fig. 2f**). Therefore, we defined regions of nuclear organisation change by their ΔPC1 score, with a cut-off of ΔPC1 ≥ |1|. This cutoff was defined by first examining ΔPC1 values within replicates of HCT116 or DKO1 Hi-C datasets, then determining the ΔPC1 values where less than 5% (2.5% either tail) of the genome would be called ‘differential’ amongst the replicates (**Supp Fig. 2g**). Called domains were merged if within 50kb.

### Hi-C compartment strength calculation

A/B compartment strengths were calculated as previously described (Stadhouders et al., 2018). Briefly, 100kb iterative corrected heat maps were generated by HiC-Pro using the hicpro2juicebox utility. 100kb bins were grouped into 50 percentile groups based on their PC1 (1^st^ eigenvector) value. Within pairwise combinations of the 50 percentile groups, average contact enrichments (obs/exp) between bins were calculated using GENOVA (v1.0.0, https://github.com/robinweide/GENOVA). Log_2_ contact enrichments were plotted as a heat saddle plot. Summarised A-A, B-B and A-B compartment strengths were calculated as the mean log_2_ contact enrichment between the top (A) or bottom (B) 20% of PC1 percentiles. The compartment strength ratio was calculated as log_2_(A-A/B-B).

### Weighted variance calculation for Repli-Seq fractions

The 6-fraction Percent-normalised Density Values (PNDVs) were used for this calculation. Briefly, PNDV values for one fraction represent the % of replication occurring within that timing fraction at any given 1kb locus. For example, the PNDV values for a locus are G1b=40, S1=38, S2=9, S3=3, S4=4, G2M=6. This means that 40% of this locus is replicated in the G1 fraction, 38% of this locus is replicated in the S1 fraction etc. This locus is biased or ‘weighted’ towards early replication timing. The sum of all fractions for any locus adds up to 100%. We made the assumption that if replication occurs evenly throughout S-phase, then each fraction from G1b to G2M should increment by 16.67% (i.e. 100%/6 fractions) to give G1=16.67, S1=33.33, S2=50.00, S3=66.67, S4=83.33, G2=100. This represents a ‘neutral’ locus that is unbiased or unweighted towards either early or late replication timing. The incremental nature of the ‘neutral’ locus informs the order of the S-phase fractions. A real locus is biased or ‘weighted’ towards early or late replication. To perform the weighted variation calculation, the PNDV values for each loci is used as the ‘weights’ against the pseudo ‘neutral’ locus, giving a measure of how the locus deviates from the ‘neutral’ locus. Formulae can be found in **Supp Fig. 4c,d** and a table of calibration tests can be found in **Supp Table 6**. The weighted variance score was discretised for visualisation purposes (**Fig. 4**) into 4 bins from 0 to 0.081 in 0.02 intervals and called var1-4.

### Single cell replication timing

#### Cell sorting and library generation using Chromium 10X Single Cell CNV Solution

We stained HCT116 and DKO1 cells using a live cell double stranded DNA dye, Vybrant DyeCycle Violet Ready Flow (Invitrogen, #R37172), according to manufacturers’ instructions. Cells were sorted (FACS Aria III) into 4-fractions: G1, Early, Mid and Late (**Supp. Fig. 10a**). Equal numbers of Early, Mid and Late cells were pooled prior to use in the Single Cell CNV system to meet the minimum recommended cell recovery number (250 cells). We aimed for 500 recovered cells. As we did not need as many G1 cells or a specific number of G1 cells, G1 cells were loaded below minimum recommended cell stock concentration and we aimed for 50 recovered cells. Single cell capture, library generation and sequencing were performed by the Garvan-Weizmann Centre for Cellular Genomics (GWCCG). Libraries were sequenced on an Illumina NovaSeq 6000 S4 flowcell (200 cycles).

#### Read mapping and filtering

Data was mapped and processed using the 10X Genomics Cell Ranger DNA (v1.1.0) software with default parameters, using the hg19/GRCh37 reference genome. Bam files generated by Cell Ranger DNA was split into individual cells/barcodes using SAMtools (Li et al., 2009) (v1.10) and filtered to remove duplicates and MAPQ < 10 reads with SAMtools, and reads overlapping hg19 blacklist (DAC) regions with BEDtools (Quinlan and Hall, 2010) (v2.26.0). Cells/barcodes with less than 1 M reads were discarded. A summary of sequencing metrics can be found in **Supp Table 7**.

#### scRepli-Seq processing

scRepli-Seq data was processed/generated based on Takahashi *et al.* (2019) with the following adjustments. Analysis was limited to autosomes.

1) Cells were filtered using median-absolute-deviation (MAD) scores, where G1 cells < 0.3 and S-phase cells > 0.4 and < 0.8 were kept for further analysis. MAD scores were calculated in non-overlapping 200kb bins. Reads were counted using *binReads* command from scCNV R package AneuFinder (Bakker et al., 2016) (v1.14.0). ‘mappable_regions.bed’ output from Cell Ranger DNA was used to generate a merged unmappable bed file of HCT116 and DKO1 for further read filtering within the *binReads* command. This applies to all further use of the *binReads* command.

2) A control dataset representing baseline copy number variations (CNVs) and mappability for S-phase cell comparison was created by merging high-quality G1 cells. CNVs in G1 cells were identified using AneuFinder *findCNVs* in 500kb bins as described by Takahashi *et al.* (2019). ‘spikiness’ and ‘bhattacharyya’ quality measures was obtained using the *clusterByQuality* command in AneuFinder. Cells were removed if spikiness >= 0.21 and bhattacharyya <= 1. Cells were also removed if deemed ‘noisy’ by Cell Ranger DNA. 48/67 HCT116 G1 and 55/84 DKO1 G1 cells passed QC and were merged for further use. To obtain a CNV baseline representative of all cells within the cell population, regions with CNV heterogeneity were removed from further analysis. Heterogeneous regions were defined as those with heterogeneity score > 0.2, calculated from the 500kb CNV data using the *karyotypeMeasures* command in AneuFinder. *karyotypeMeasures* was modified to output heterogeneity score per bin. Heterogeneous regions from HCT116 and DKO1 were merged and removed from all further analyses.

3) Single cell data was normalised against the merged G1 control data. G1 and S-phase cell reads were binned in non-overlapping 80kb bins and in 200kb bins at 40kb sliding intervals using AneuFinder *binReads*. Read counts were normalised using the *correctMappability* command from AneuFinder (copied from v1.5.0), using the merged G1 data as reference control.

4) Single cell RT scores were generated from the mappability-corrected 200kb bin-40kb sliding window data as described in Takahashi et al. (2019). Single cell replication timing profiles are similar to whole population Repli-Seq timing profile (**Supp. Fig 10b**). Single cell RT scores were used to generate Pearson’s correlation matrix, hierarchical clustering (ward.d2) and tSNE plot as described in Takahashi *et al.* (2019). At this point, any S-phase cells that cluster with G1 in the hierarchical clustering or in the tSNE plot were removed from further analysis. 322/646 HCT116 S and 208/582 DKO1 S cells were used for further analysis.

5) Mappability-corrected 80kb bin data was binarized using the *findCNVs* command in AneuFinder. We used the parameters specified in Takahashi *et al.* (2019): method=“HMM”, max.iter=3000, states=c(“zero-inflation”, “0-somy”, “1-somy”, “2-somy”), eps=0.01, most.frequent.state=(“1-somy” or “2-somy”). The *findCNVs* command outputs a 2-state HMM model were states ‘1’ and ‘2’ indicates un-replicated and replicated, respectively. Within the *findCNVs* command, Takahashi *et al.* (2019) specifies whether the most common state is 1-somy (unreplicated) or 2-somy (replicated) depending on FACS gating to reduce HMM calling ambiguity, i.e. a Early cell would have majority state ‘1’ and a Late cell would have majority state ‘2’. However, as Early, Mid and Late S-phase cells were pooled here, we could not directly assign whether 1-somy or 2-somy was the most common state. Therefore, we generated two HMM models per cell, specifying either 1-somy or 2-somy as the most common state. For most cells, the two HMM models had little to no difference in binarisation. These tended to be Mid-S cells (**Supp. Fig. 10c**). We determined that a cell had ‘evenHMM’ if the absolute differences in bin numbers between 1-somy state ‘1’ and 2-somy state ‘1’ is less than 1,500 bins, and the same for state ‘2’. A cell was ‘unevenHMM’ if the absolute differences were greater than 1,500 bins for both states. The cutoff of 1,500 bins was conservatively chosen to separate the two groups based on the distribution of bin number differences between 1-somy and 2-somy calls of the same state. This distribution was bimodal, with one group centred around 0 (‘evenHMM cells) and the other peak centred around 15,000-20,000 (‘unevenHMM’ cells) (**Supp. Fig. 10d**). ‘unevenHMM’ cells were then assigned as Early or Late through visual comparison of the HMM bed file against the earliest and latest ‘evenHMM’ cells. For final HMM calls, we used the 2-somy calls for ‘evenHMM’ cells, the 1-somy calls for ‘unevenHMM’ cells that were assigned as Early and the 2-somy calls for ‘unevenHMM’ cells that were assigned as Late. Binarized 80kb data was then used to generate the % replication score per cell, single cell RT value per bin, cell-to-cell variability scores and RT sigmoid modelling (gain and M values) as described by Takahashi *et al.* (2019). For calculation of the slope of the sigmoid model (gain) instead of using an estimated 10-h S-phase time window, we used the % replication score that the 10-h time window was based on in Takahashi *et al.* (2019). Thus our gain values are on a different scale to those in Takahashi *et al.* (2019). The M-value is the x-intercept at the sigmoid’s midpoint and represents when 50% of the cell population has replicated that locus.

### Identifying biphasically replicating loci

Loci were called as biphasic if their weighted variance score was ≥ 0.081. Calibration tests of the weighted variance score showed that a score of ~0.081 is achieved for a locus with even replication timing across all 6 fractions (**Supp Table 6**, ‘even’), and a score above ~0.081 represents loci where there were high PNDV values in non-adjacent fractions separated by in-between fractions of low PNDV values (**Supp Table 6**). The cut-off of 0.081 does miss some biphasic regions with smaller separations between high PNDV fractions, hence, the use of the Hansen score described below. Biphasic loci were also identified according to Hansen *et al.* (2010). Briefly, the 6-fraction PNDV scores were reduced to 5 fractions by pairwise addition of adjacent fractions (G1b+S1, S1+S2, S2+S3, S3+S4, S4+G2M). A 1kb locus was deemed biphasic if more than 40% of the 5-fraction score was in non-adjacent fractions. For example, a locus is biphasic if ≥ 40% of the score is in G1+S1 and ≥40% is in S2+S3, S3+S4 or S4+G2, with S1+S2 < 40%.

Biphasic loci are defined as *maintained* where at least one HCT116 replicate and one DKO1 replicate are biphasic. Biphasic loci are defined as *gained* where no HCT116 replicates are biphasic and at least one DKO1 replicate is biphasic. Biphasic loci are defined as *lost* where at least one HCT116 replicate is biphasic and no DKO1 replicates are biphasic.

### Allelic replication timing

#### Nanopore sequencing, base calling, alignment, variant calling and phasing

HCT116 and DKO1 DNA (1 μg) was sheared using the Covaris g-TUBE spun at 3400 x g in 2 × 60 sec spins. Sheared DNA was prepared for Nanopore sequencing using the Ligation 1D kit (SQK-LSK109) according to manufacturer’s instructions. Each cell line was sequenced on one PromethION flow cell. Reads were base called with Guppy v3.3.0 on a GPU-enabled Sun Grid Engine high performance computing server (parameters “--chunks_per_runner 1500 --gpu_runners_per_device 1 --cpu_threads_per_caller 4 -x “cuda:0 cuda:1 cuda:2 cuda:3” -r” and configuration “dna_r9.4.1_450bps_hac_prom.cfg”). Base called reads (fastq) were aligned to hg19 using minimap2 (Li, 2018) (v.2.17-r943-dirty) with parameters “-ax map-ont”. Mapped reads were sorted and indexed with SAMtools (v1.9). Variants were called and phased with medaka_variant (v0.11.4, https://nanoporetech.github.io/medaka) with the options “-t 36 -s r941_prom_high_g330 -m r941_prom_high_g330 -p -b 100”.

#### Variant filtering, haplotype mapping and Repli-Seq processing

Medaka variants were filtered as follows using bcftools (v1.9): i) occurs in both HCT116 and DKO1 datasets; ii) quality score above 20; iii) within each dataset, only phased heterozygote single nucleotide variants (SNVs) were used. hg19 reference genome fasta files were generated for each haplotype per cell line using bcftools consensus (parameters “-H 1pIu” and “-H 2pIu”). Repli-Seq fractions were mapped to each haplotype reference genome using bowtie (Langmead et al., 2009) (v1.1.0, parameters “-v 0 -m 1 --tryhard --best --strata --time --trim5 6”). Mapped bam files were filtered for reads that overlapped phased SNVs using SAMtools view (v1.9, option “-L”). Weighted average scores were calculated from filtered bam files as described above with the following modifications: i) Reads were counted in 50 kb sliding windows at 1 kb intervals. The 50 kb sliding windows were modified so that only reads within each haplotype block were counted for each block; ii) The low coverage threshold for 50kb windows was set to at least 5 reads per fraction per 50kb loci.

#### Calling allelic differentially timed regions

Allelically replicating regions were called if there was a WA difference of less than 10 between replicates and more than 30 between alleles. Due to the sparseness of allelically mapped regions, allelic regions were merged if within 1 Mb of each other. The 1 Mb merged region was used to identify overlapping genes.

#### Imprinted gene and cancer-related gene annotation

A list of human imprinted genes was obtained from www.geneimprint.com (Luedi et al., 2007). Genes located in allele-specific replication regions were also checked against the Candidate Cancer Gene Database (**Supp Table 1**) (http://ccgd-starrlab.oit.umn.edu) (Abbott et al., 2015).

### Profile plots

We used SeqPlots (Stempor and Ahringer, 2016) to calculate average scores over regions of interest, then used ggplots (Wickham, 2016) to plot the average scores across all regions for each bin with standard error and confidence intervals.

### ChIP-seq processing

ChIP-seq datasets were processed as previously described (Bert et al., 2013; Taberlay et al., 2014). Briefly, ChIP-seq reads were aligned to hg19 using bowtie (Langmead et al., 2009) (v1.1.0) allowing up to 3 mismatches, discarding ambiguous and clonal reads. All histone ChIP-seq peaks were called using PeakRanger (Feng et al., 2011) (v1.16). For the distribution of histone mark occupancy across replication timing, consensus peaks were used where replicates were available. Broad domains of histone mark enrichment, H3K4me3 and H3K9me3, were processed with Enriched Domain Detector (EDD) (Lund et al., 2014) (settings: required_fraction_of_informative_bins = 0.9, p_hat_CI_method = normal). Replicate-shared domain for H3K9me3 were called using the ‘intersect’ function (R, GenomicRanges (Lawrence et al., 2013)), before calling regions of maintenance (‘intersect’ between HCT116 and DKO1), gain and loss (‘setdiff’ between HCT116 and DKO1). H3K4me3 domains that exist only in DKO1 (replicate merged) were intersected with H3K9me3 domain regions and used for further analyses.

### ChromHMM analysis

15-state ChromHMM tracks for HCT116 and DKO1 were called based on the Roadmap Epigenomics 15-state chromHMM model (Roadmap Epigenomics et al., 2015) using the chromHMM program (v1.10) (Ernst and Kellis, 2012). ChIP-seq data was prepared for segmentation by first using ‘bamToBed’, followed by ‘BinarizeBed’. Replicates were pooled at the bamToBed stage. The Roadmap 15-state model parameters were then applied to produce 15-state segmentations for HCT116 and DKO1. Analysis of ChromHMM change-of-state is based on Fiziev et al. (2017) (Fiziev et al., 2017). To calculate ChromHMM state change enrichment scores, we divided the number of observed state changes by the number of expected changes as outputted by the chisq.test in R. Two-sided p-values were calculated from the chisq.test standard residuals (similar to the z-score) and FDR corrected. To control for reciprocal state changes (i.e. Het to TssA versus TssA to Het in the direction of HCT116 to DKO1), the enrichment scores of Het-TssA was divided by the enrichment score of TssA-Het. A count of 1 was added to both observed and expected to avoid divisions by 0. Only transitions where both scores were significant are shown.

### RNA-seq data generation and processing

Total RNA in triplicates (different passages) was extracted from cultured cells using TRIzol (Life Technologies, #15596018). Libraries were constructed with the Illumina TruSeq Stranded mRNA library preparation kit (Illumina, #RS-122-2102) and sequenced on the Illumina HiSeq X Ten. Paired-end reads were processed as previously described (Du et al., 2019) using Trim Galore (v0.4.0, parameter settings: --fastqc --paired --retain_unpaired --length 16) and STAR (Dobin et al., 2013) (v2.5.3a, parameter settings: --quantMode TranscriptomeSAM) for mapping reads to the hg19 human transcriptome build (GENCODE 19 (Harrow et al., 2012)). Mapped reads where counted into genes using rsem (v1.2.21) (Li and Dewey, 2011). TMM normalization was applied using edgeR (v3.12.1) (Robinson et al., 2010). Fold changes (FC) were computed as the log2 ratio of normalized reads per gene using edgeR. Genes with fold change ± 1.5 and FDR < 0.01 were considered as significantly altered. Promoters were defined as the region from −2000 bp to +100 bp around the transcriptional start site.

### Statistical tests

For genomic interval overlaps and genomic rearrangement overlaps, a modified LOLA (Sheffield and Bock, 2016) package was used to perform a two-sided log-odds ratio test which reports significance using “BH” FDR values. The Mann-Whitney-Wilcoxon test was used for 2-group non-parametric comparisons, and the one-tailed test was used where a directional difference between the groups was of interest. Unless otherwise stated, statistical tests were two-sided.

### External data

ChIP-seq datasets were downloaded from GSE58638, performed by Lay *et al.* (2015).

### Data availability

Repli-Seq, Hi-C, WGBS, single cell Repli-Seq, Nanopore sequencing and RNA-seq are available from the NCBI Gene Expression Omnibus (GEO) under accession number GSE158011.

