## Supplemental Figures for "DNA methylation is required to maintain DNA replication timing precision and 3D genome integrity"

Du_et_al_Supp_Figure_1

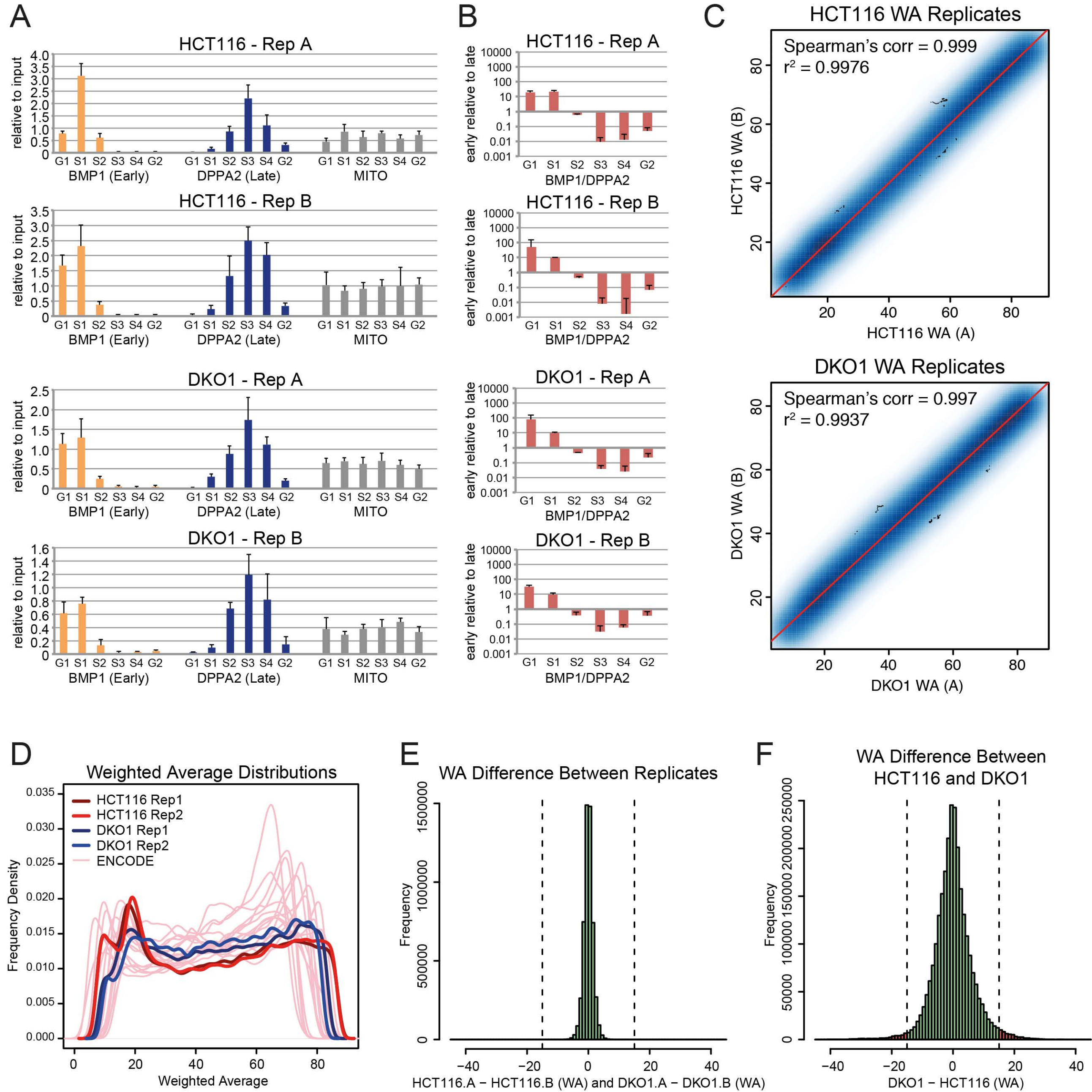

Du_et_al_Supp_Figure_2

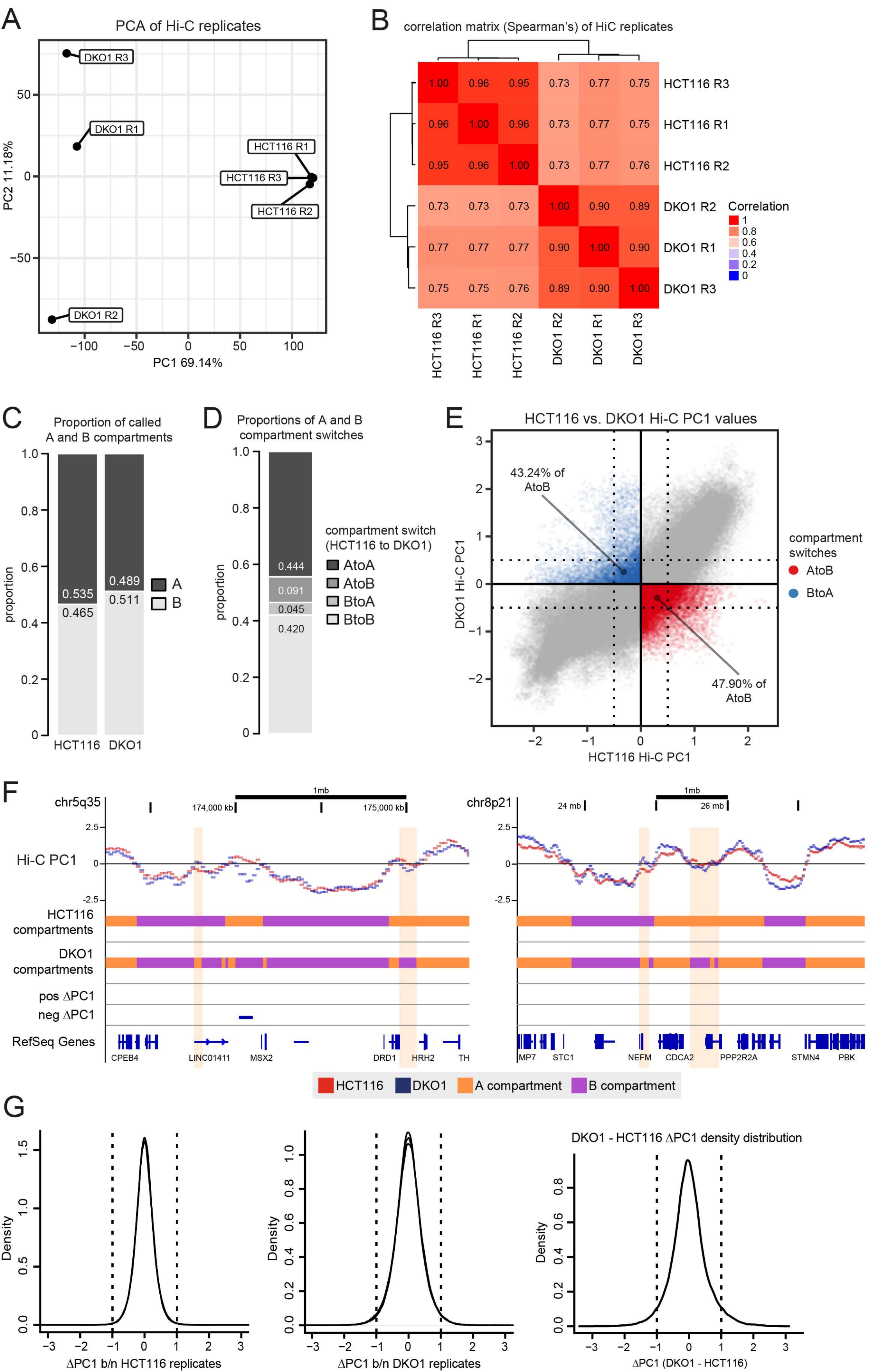

Du_et_al_Supp_Figure_3

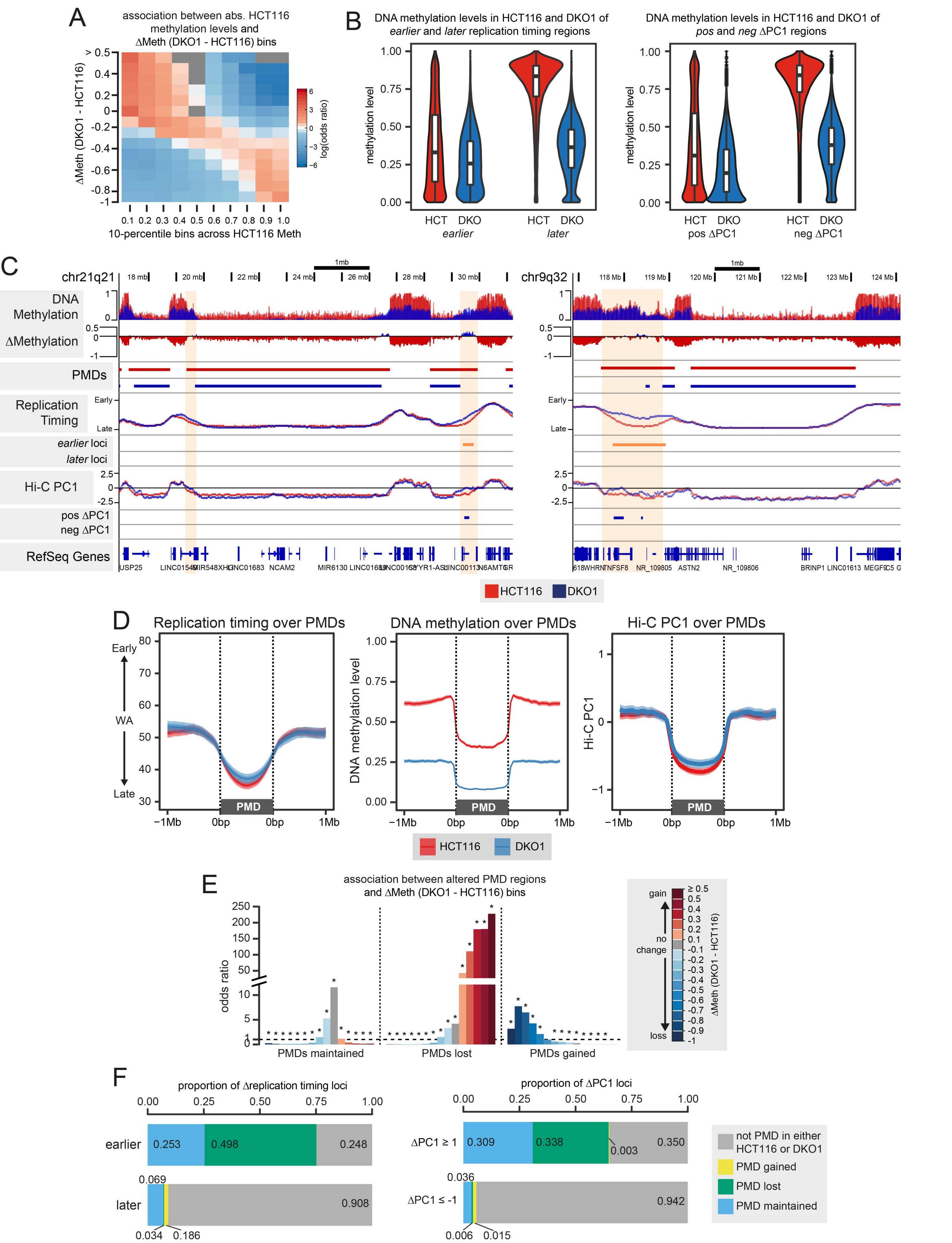

Du_et_al_Supp_Figure_4

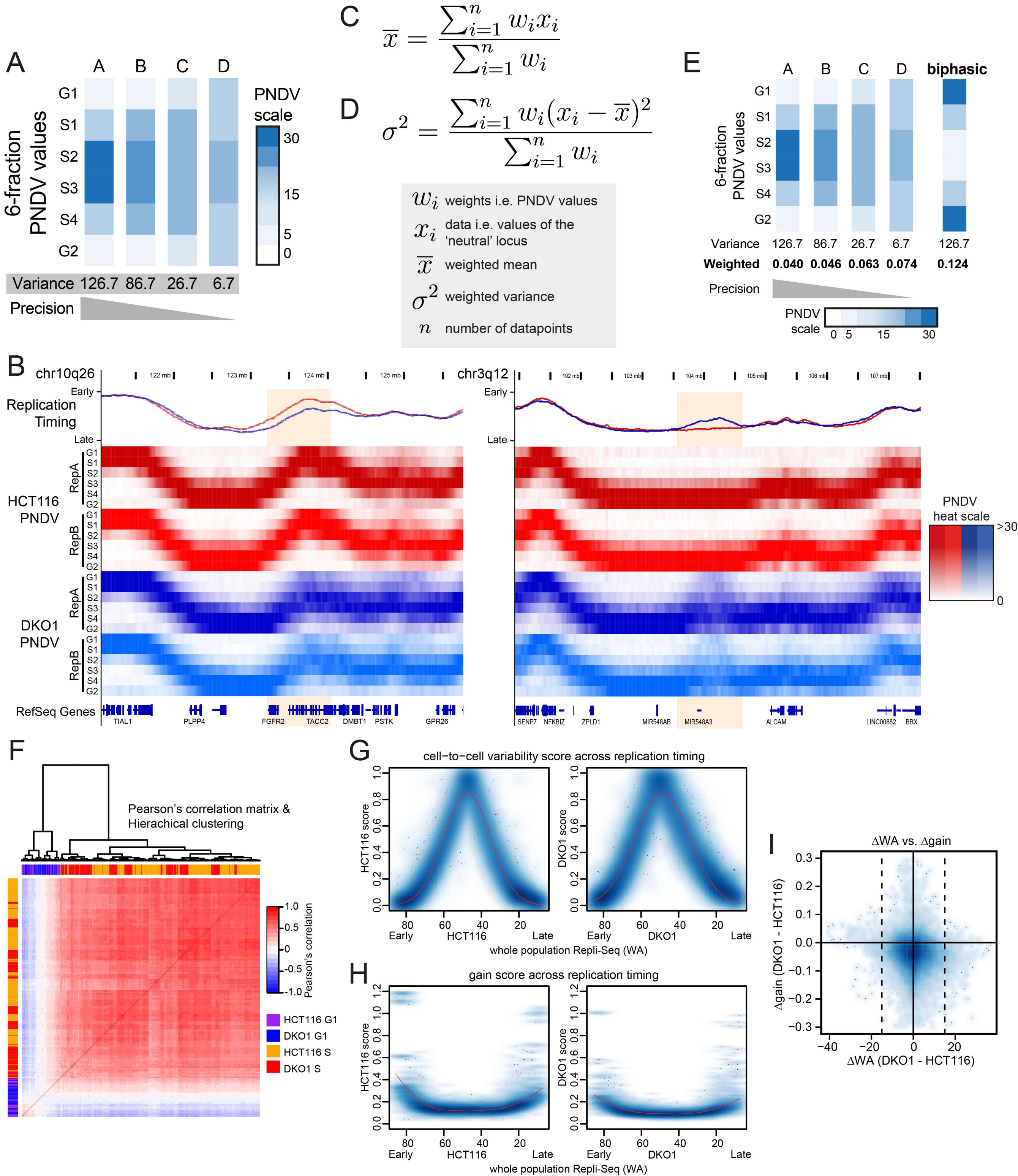

Du_et_al_Supp_Figure_5

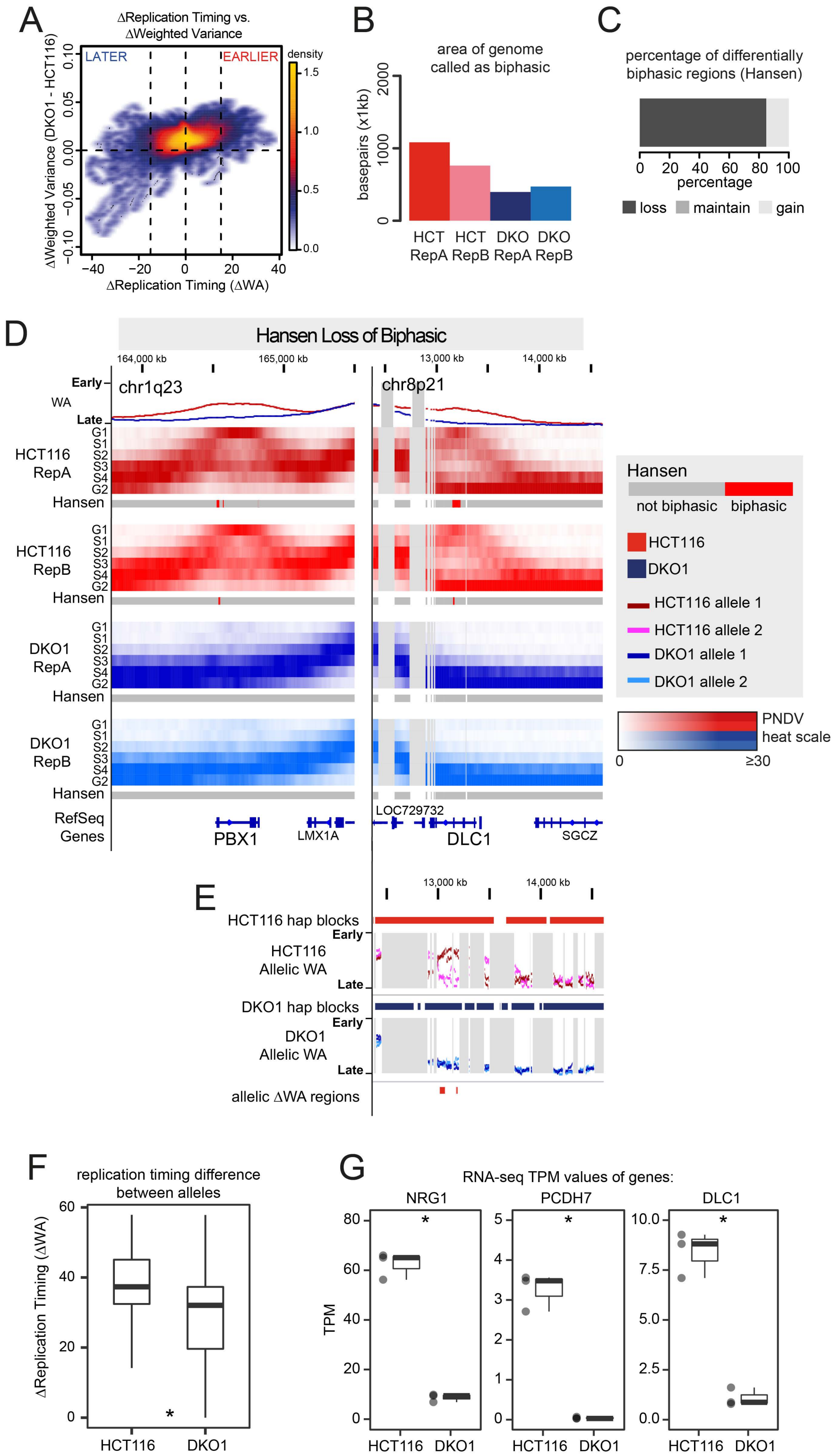

### Du_et_al_Supp_Figure_6

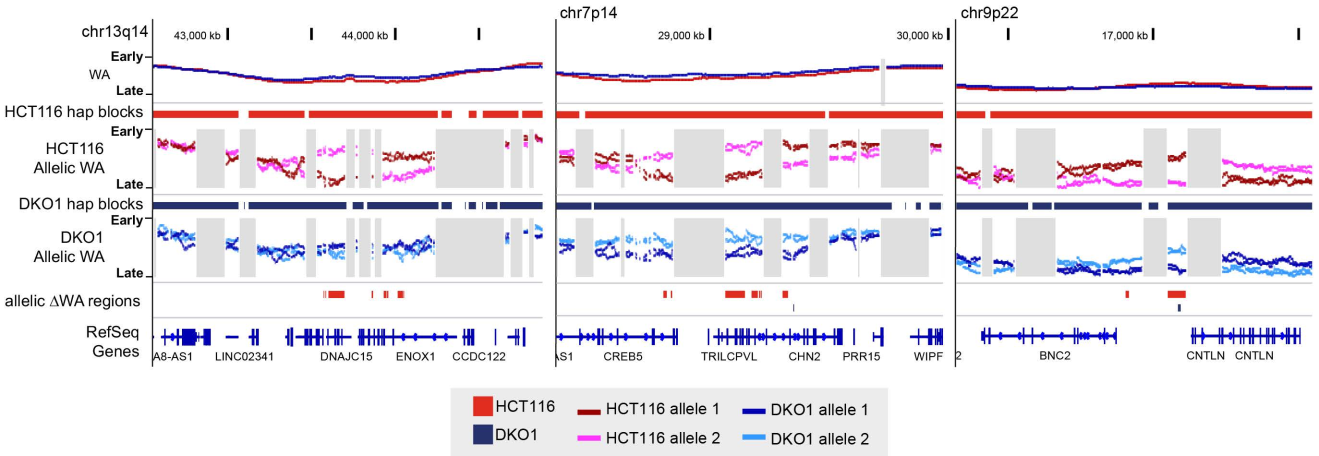

Du_et_al_Supp_Figure_7

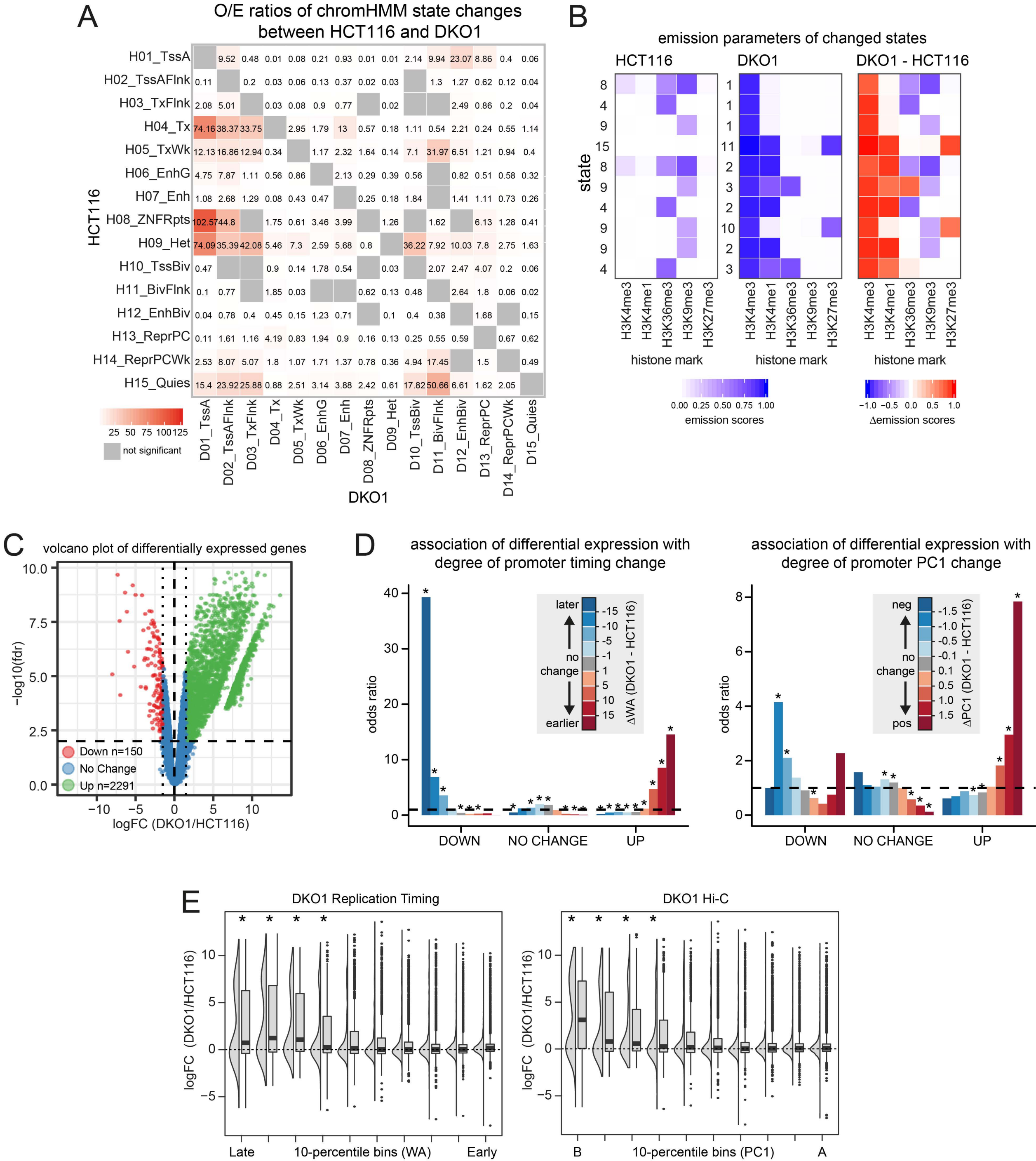

Du_et_al_Supp_Figure_8

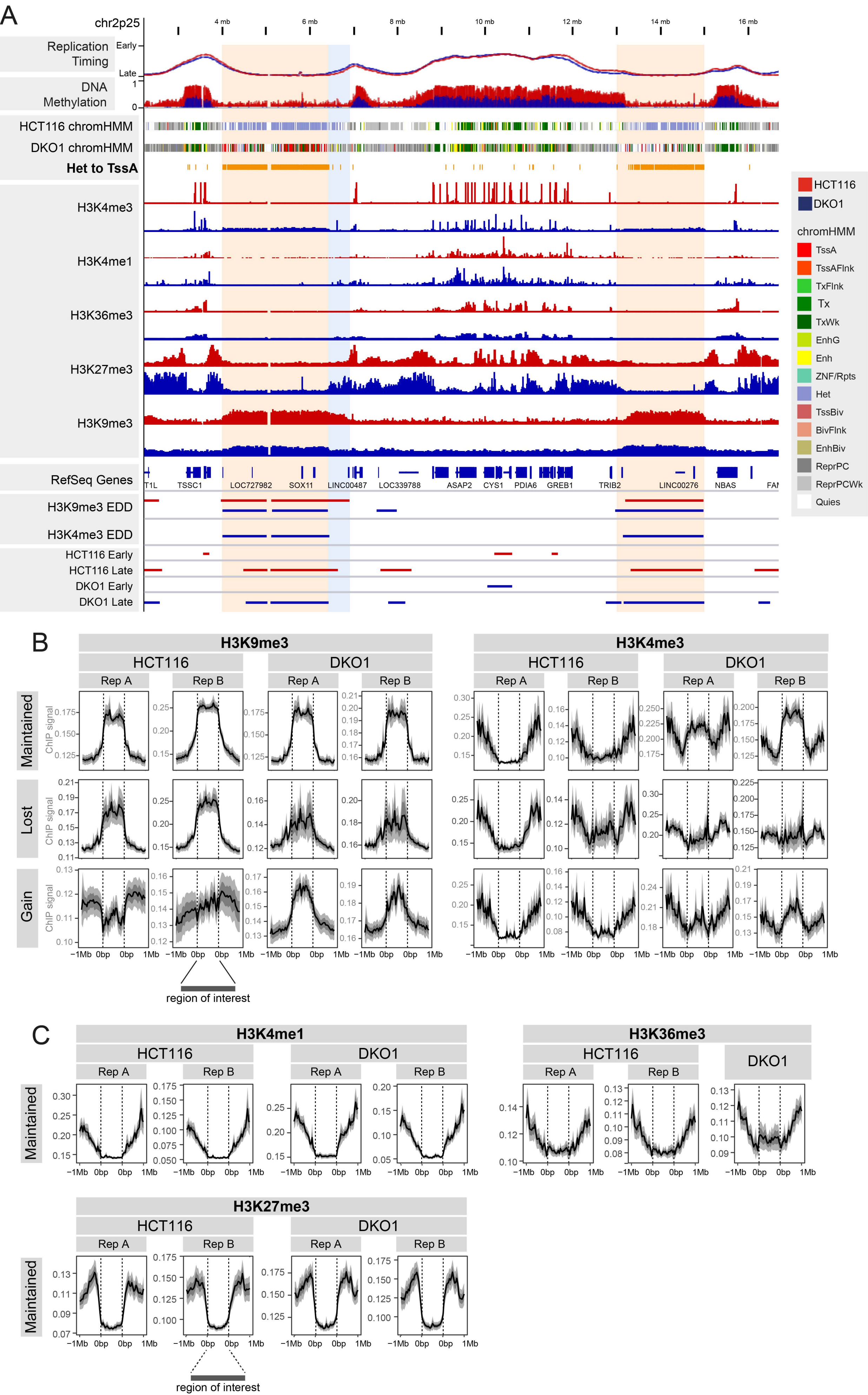

Du_et_al_Supp_Fig_9

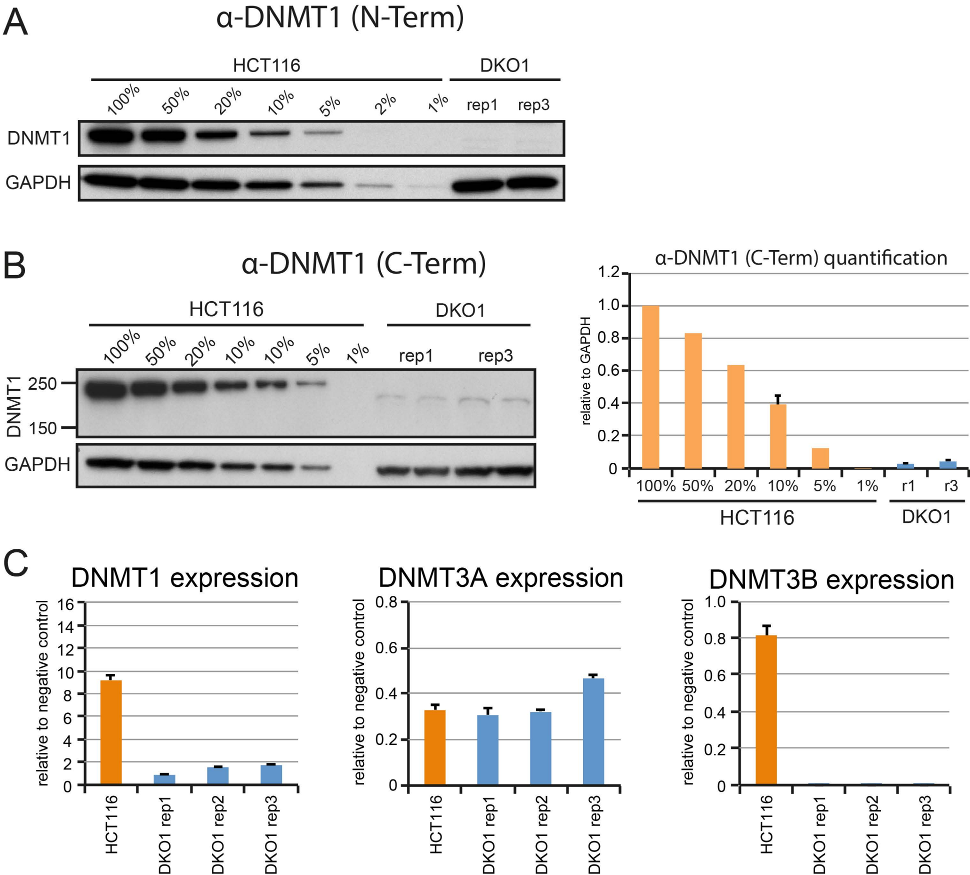

Du_et_al_Supp_Figure_10

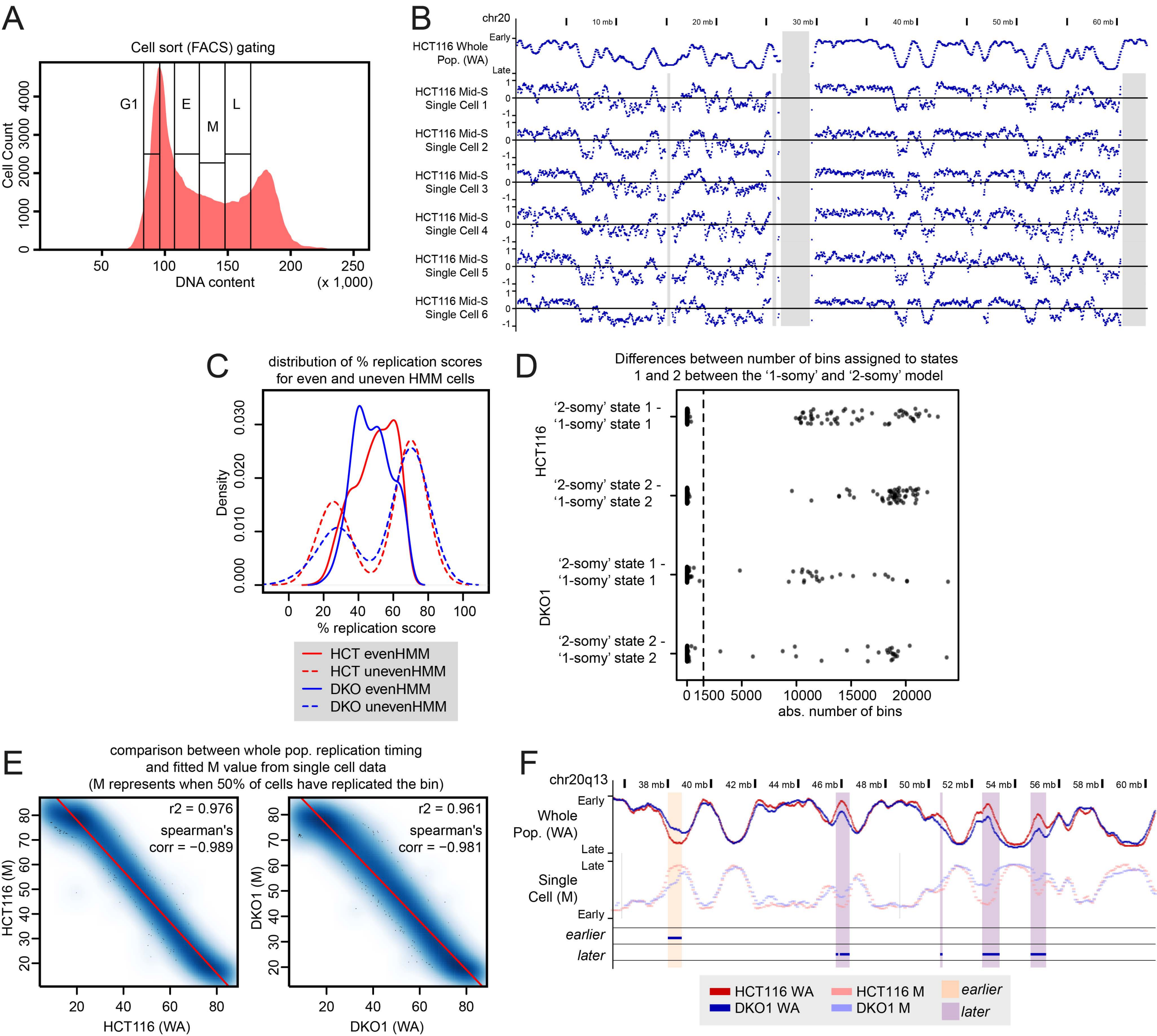

#### Supplemental Figure Legends

**Supp Figure 1: Repli-Seq of HCT116 and DKO1 cells.**

**A** Validation of S-phase sorting. qRT-PCR of known early (BMP1) and late (DPPA2) loci were carried out on BrdU-labelled DNA extracted from sorted HCT116 and DKO1 samples. A locus on the mitochondrial genome is used as an S-phase independent control, as the mitochondrial genome replicates independently of the nuclear genome. Y-axis shows enrichment of loci against input control. Error bars are SD. **B** Relative qRT-PCR enrichments of the early loci (BMP1) against the late loci (DPPA2). The y-axis uses a $\log_{10}$ scale. Error bars are SD. **C** Replicates of each cell line show high Spearman's correlation and $r^2$ scores. **D** The distribution of HCT116 (red) and DKO1 (blue) WA values are comparable to the WA distributions of twelve ENCODE Repli-Seq datasets (pink). The distribution of WA differences between replicates (**E**) is compared to the distribution of WA differences between HCT116 and DKO1 (**F**). Dotted lines indicate a $|\Delta WA| > 15$. Values that fall outside this range signify loci that have changed replication timing.

**Supp Figure 2: Hi-C of HCT116 and DKO1 cells.**

**A** PCA plot of Hi-C replicates for HCT116 and DKO1. **B** Correlation matrix of Hi-C PC1 values between cell lines and replicates. Values are Spearman's correlation. **C** Proportion of the genome called as either A- or B-compartments in HCT116 and DKO1. **D** Proportion of the genome showing A/B-compartment maintenance or switching between HCT116 and DKO1. **E** Scatterplot of HCT116 PC1 values against DKO1 PC1 values, showing that the majority of A-B compartment switches are close to PC1 values of 0 (PC1 < |0.5|, dotted lines). **F** Representative examples of A-B compartment switches with little difference in PC1 values between HCT116 and DKO1. **G** Distribution of PC1 differences between replicates of HCT116 and DKO1, and between HCT116 and DKO1. Dotted lines indicate a $|\Delta PC1| > 1$, where less than 5% of the genome would be called 'differential' amongst the replicates.

**Supp Figure 3: *Earlier* replication timing and PC1 changes corresponds to loss of partially methylated domains (PMDs) in DKO1.**

**A** Association between 10-percentile bins of absolute HCT116 methylation levels and bins of DNA methylation change. Only significant associations (FDR < 0.05, Fisher's exact test) are shown, non-significant associations are shaded in grey. **B** Methylation levels of *earlier* and *later* regions, and domains of large PC1 change ($\Delta PC1 >= |1|$) in HCT116 and DKO1. **C** Representative examples of lost PMD regions, particularly at PMD boundaries. All HCT116 datasets are shown in red and all DKO1 datasets are shown in blue. Beige highlighting indicates regions of PMD loss from HCT116 to DKO1. **D** Profile plots of average HCT116 and DKO1 replication timing, DNA methylation and Hi-C PC1 across PMDs of each cell line. Plots show an average line with width of shading indicating confidence intervals. **E** Association between altered PMD regions and bins of DNA methylation change. Only significant associations (FDR < 0.05, Fisher's exact test) are shown, non-significant associations are shaded in grey. **F** Contribution of altered PMD regions to domains of replication timing change ($\Delta WA > |15|$) and domains of Hi-C PC1 change ($\Delta PC1 >= |1|$).

**Supp Figure 4: Global DNA methylation loss increases intra-population heterogeneity of DNA replication.**

**A** Example loci from A to D, showing progressively lower variance scores and progressively lower precision of the 6-fraction PNDV signal spread (percentage normalised density values, See Methods). **B** Representative examples where the variance/spread of DNA replication across DKO1 (blue) PNDV fractions is higher than HCT116 (red) (beige shaded regions). All 6 PNDV fractions of each Repli-Seq datasets are shown in the order of G1, S1, S2, S3, S4, and G2 from top to bottom. PNDV value for each fraction is indicated by a heat colour scale. **C** Formula for calculating the weighted mean. **D** Formula for calculating the weighted variance using the weighted mean. As the weights ($w_i$) are the percent normalised density values (PNDVs), the sum of weights adds up to 100. The number of data points ($n$) is 6 for the 6 fractions. **E** The same example loci from **A,** except now with their weighted variance scores, showing that loci with *higher* weighted variance scores have lower precision. An even higher weighted variance score indicates biphasic loci (See Methods). **F** Pearson's correlation matrix and clustering of single cell Repli-Seq data. **G** Heat scatterplot of cell-to-cell variability score across whole population replication timing (WA). Red line indicates Gaussian fitting of the cell-to-cell variability data calculated as described in Takahashi *et al.* (2019). **H** Heat scatterplot of gain across whole population replication timing (WA). Red dotted line indicates loess curve fitting of data. **I** Heat scatterplot of DKO1-HCT116 difference in gain against change in replication timing. Vertical dotted lines indicate ΔWA cutoffs of -15 and 15.

**Supp Figure 5: Loss of DNA methylation causes loss of biphasic DNA replication.**

**A** Change in replication timing (DKO1 – HCT116) against the change in weighted variance (DKO1 – HCT116). Density is indicated by colour scale. **B** Area of the genome in kilobases (Kb) called as ‘biphasic’ using the Hansen method. **C** Percentage of biphasic replication regions that are lost, maintained or gained from HCT116 to DKO1 as calculated using the Hansen method. **D** Examples of regions that changed biphasic status between HCT116 and DKO1 as calculated using the Hansen method. All 6 PNDV fractions of each Repli-Seq datasets are shown in the order of G1, S1, S2, S3, S4, and G2 from top to bottom. HCT116 datasets are shown in red and DKO1 datasets are shown in blue. The weighted average replication timing score is shown at the top of each example. Colours indicating Hansen categorisation is shown below each lot of 6 fractions. PNDV value for each fraction is indicated by a heat colour scale. **E** Allelically separated replication timing (WA) scores of the same regions in **D**. Haplotype blocks are shown at the top of each example. Grey shading indicates either no data or a break between haplotype blocks. **F** Boxplots of replication timing difference (ΔWA) between alleles in HCT116 and DKO1. Asterisk indicate that ΔWA in DKO1 is less than in HCT116 ($p < 0.05$, one-tailed Mann-Whitney-Wilcoxon). **G** RNA-seq TPM values of HCT116 and DKO1 for *NRG1*, *PCDH7* and *DLC1*. Asterisks indicate the gene is significantly differentially expressed (FDR < 0.01, logFC > |1.5|).

**Supp Figure 6: Examples of allelic replication timing in HCT116 that is reduced or lost in DKO1.**

Non-allelically separated replication timing scores (WA) are shown in the top track and allelically separated replication timing scores for HCT116 and DKO1 are shown in the second and third tracks. For the allelic tracks, haplotype blocks are shown at the above the WA scores. Grey shading indicates either no data or a break between haplotype blocks. Allele-specific regions called in either HCT116 (red) or DKO1 (blue) are shown in the fourth track.

**Supp Figure 7: Chromatin and expression changes after DNA methylation loss corresponds to replication timing change and genome reorganisation**

**A** Normalised chi-squared observed/expected ratios (O/E) for genome-wide chromHMM chromatin state changes between HCT116 and DKO1. Only significant changes are shown (FDR < 0.05, chi-squared test). **B** Emission parameters of the top 10 states that change between HCT116 and DKO1, ranked from highest to lowest O/E ratios. The emission intensities indicate the probability of the histone mark to be found in that state. The 'Δemission scores' heatmap indicate which marks have changed between the paired states. **C** Volcano plot of differential expression between HCT116 and DKO1. All testable genes are shown. **D** Association between differential gene expression categories and bins of promoter replication timing change or promoter Hi-C PC1 change. Asterisks indicate significant associations (Fisher's exact test, FDR < 0.05). **E** Boxplots and density plots of differential expression (logFC) in each of 10-bins of replication timing (WA) or Hi-C PC1 values of DKO1. Asterisks indicate significance in one-tailed Mann-Whitney-Wilcoxon test of each bin against genome-wide differential expression, where the alternative 'greater'.

**Supp Figure 8: Maintained H3K9me3 regions between HCT116 and DKO1 gain H3K4me3 ChIP-seq signal in DKO1.**

**A** Representative example where H3K9me3 is maintained from HCT116 to DKO1 and H3K4me3 is gained in DKO1. All HCT116 datasets are shown in red and all DKO1 datasets are shown in blue. The 'Het to TssA' track shows loci of chromatin state transition. Beige shading indicates where H3K9me3 domains are maintained from HCT116 to DKO1. Blue shading indicates where H3K9me3 domains are lost from HCT116 to DKO1. **B** H3K9me3 and H3K4me3 ChIP-seq profile plots over H3K9me3 regions that are maintained, lost, or gained from HCT116 to DKO1. **C** H3K4me1, H3K36me3 and H3K27me3 ChIP-seq profile plots over H3K9me3 regions that are maintained from HCT116 to DKO1. For **B** and **C**, profile plots and heatmaps are shown for the region of interest and extended by 1Mb on either side.

**Supp Figure 9: Validating knockout status of DKO1 cells.**

**A** Western blot of DNMT1 using an N-terminal antibody targeting the deleted region. **B** Western blot of DNMT1 using a C-terminal antibody targeting the truncated protein. DNMT1 quantitation relative to loading control GAPDH is shown on the right. **C** Expression levels of *DNMT1*, *DNMT3A* and *DNMT3B* in HCT116 and replicates (different passages) of DKO1. Error bars are SD. For Western blots, serial dilution of HCT116 protein lysate is used as a quantitative comparison for DKO1 protein levels. Western blot quantitation was performed in ImageJ and where replicates were available, error bars are calculated as SEM.

**Supp Figure 10: Processing single cell Repli-Seq.**

**A** Cells were sorted into 4 gates: G1, Early (E), Mid (M) and Late (L). **B** Representative example of the similarity between whole population Repli-Seq profile (HCT116) and single cell Repli-Seq profiles. Six mid-S HCT116 single cell profiles are shown. **C** Single cell % replication score for 'even' and 'uneven' HMM cells. **D** Jitter plot showing distribution of differences in number of bins assigned to state 1 or

2 between ‘1-somy’ and ‘2-somy’ HMM models. Vertical dotted line represents cut-off of 1500.
